# Linking biosurface reactivity and photosynthesis to investigate *Sphagnum* mosses as peatland engineers

**DOI:** 10.64898/2025.12.10.693362

**Authors:** Anna Di Palma, Emanuele Pallozzi, Aridane G. González, Oleg S. Pokrovsky, Ralf Reski, Janice Glime, Norbert Kavasi, Carlo Calfapietra

## Abstract

- *Sphagnum* mosses regulate peatland carbon cycling, hydrology, nutrient dynamics, and structure. We investigated the link between *Sphagnum* surface chemistry and photosynthetic efficiency, and the combined effects of pH and waterlogging on photosynthesis, key mechanisms for peatland functioning but still poorly understood.
- We quantified biosorption potential and functional groups in 20 field collected and in 4 axenically cloned *Sphagnum* species via titrations. Gas exchange and chlorophyll fluorescence measurements were performed in 5 representative species under submerged conditions across a pH gradient.
- The 20 *Sphagnum* species share a basic surface chemistry, but they differ in surface charge and functional group abundance, with an overall greater biosorption potential in the subgenera *Acutifolia* and *Sphagnum*. Photosynthesis is species- and pH-dependent, influenced by surface functional-group composition, with phosphoryl and carboxyl groups enhancing CO_2_ assimilation. *Sphagnum palustre* emerges as a generalist, stress-tolerant species, maintaining stable photosynthesis and photoprotection across pH ranges. *In-vitro* cultivation reduces surface reactivity and interspecific differentiation but preserves fundamental *Sphagnum* surface chemistry.
- These results highlight the interplay between chemical reactivity, photosynthetic performance, and ecological adaptation in *Sphagnum*. *Sphagnum* chemical traits and photosynthetic plasticity enable survival under variable pH, waterlogging, and nutrient conditions, with clones providing reliable models for research and applications.

## Introduction

Peat mosses from the genus *Sphagnum* have experienced 350 million years of evolution separate from all other mosses (Lueth & Reski, 2023) and are unique among bryophytes in terms of ecological role, particularly in the context of carbon cycling. These peat-forming species dominate northern peatlands, which store approximately 30 % of global soil carbon as partially decomposed organic matter, known as peat (Liu *et al*., 2024; Minasny *et al*., 2024). Through their high cation exchange capacity, *Sphagnum* mosses create an acidic and nutrient-poor environment, fostering waterlogged, anoxic, and cold conditions that greatly slow decomposition and promote peat accumulation (Clymo & Hayward, 1982; Turetsky *et al*., 2012; Rydin & Jeglum, 2013). As a result, peatlands function as the most effective long-term terrestrial carbon sinks (Limpens *et al*., 2008; Page *et al*., 2011; Yu *et al*., 2021).

However, these ecosystems are very vulnerable to climate changes and anthropogenic disturbances, such as warming, drought, land use transformation, and overexploitation of peat as fuel resource (Charman *et al*., 2013; Hopple *et al*., 2020; UNEP, 2022; Ofiti *et al*., 2023). When destabilized, peat decomposition accelerates, releasing CO_2_ and CH_4_, and risking a transition from carbon sink to carbon source (Bridgham *et al*., 2013; Joosten, 2015; Bragazza *et al*., 2016; Hopple *et al*., 2020). Such feedback can amplify global climate warming (Rafat *et al*., 2021), underscoring the urgency of peatland conservation and restoration (Andersen *et al*., 2017; Humpenöder *et al*., 2020; Doelman *et al*., 2023). *Sphagnum*-based approaches, including *Sphagnum* farming and axenic cultivation (Gaudig *et al*., 2014; Parson *et al*., 2025), are increasingly recognized as promising nature-based solutions, offering a sustainable alternative to conventional peat extraction while maintaining peatland ecosystem functions (Temmink *et al*., 2024).

In addition to carbon storage, *Sphagnum* mosses are known for their biosorption capabilities, owing to the abundance of surface functional groups that bind metal ions and other pollutants (González & Pokrovsky, 2014; González *et al*., 2017). These properties have made them effective tools in environmental monitoring, including atmospheric pollution assessments (Di Palma *et al.,* 2017, 2022), and historical reconstructions of atmospheric depositions and climate changes via peat core profiles (e.g., Shotyk *et al.,* 2015; Davies *et al.,* 2018). By releasing allelochemicals and acting as nutrient filters, *Sphagnum* mosses are ecological engineers, responsible for regulating plant succession and habitat selection in peatlands, thereby maintaining and shaping these ecosystems (Rydin *et al*., 2006; Hamard *et al*., 2019). Moreover, they have multiple adaptations highly specialized for water retention (*e.g.,* hyaline cells; Clymo & Hayward, 1982; van de Koot *et al*., 2024), contributing significantly to the peatland hydrological cycle and overall water balance (Gauthier *et al*., 2022).

Despite this central role, the surface chemistry of *Sphagnum* and its link to photosynthetic physiology remain poorly understood. While previous research has focused on peatland hydrology and carbon fluxes, the interplay between proton exchange capacity, pH regulation, and photosynthetic efficiency, processes that are crucial for both carbon fixation and pollutant uptake by *Sphagnum*, has not been systematically addressed. This gap is particularly relevant given the lack of stomata in *Sphagnum* gametophytes (McAdam *et al*., 2021), which makes CO_2_ assimilation highly sensitive to water availability and pH. Although drought effects have been studied (Schipperges & Rydin, 1998; Harris, 2008; Jassey & Signarbieux, 2019; Keane *et al*., 2025), the combined influence of pH and water submersion on moss photosynthesis remains largely unexplored.

Here, we present the first integrated study combining acid–base titration with photosynthetic measurements to elucidate the surface reactivity of *Sphagnum* mosses, and their physiological response under simulated peatland water-saturation conditions. Photosynthesis is the process used by *Sphagnum* to fix atmospheric CO_2_, and hence to stock it as peat (Serk *et al*., 2021). Surface groups are involved in bioadsorption, a physical-chemical process considered the first step of the accumulation of sorbates (pollutants included) into the cells and responsible for proton displacement and ion exchanges (including dissolved CO_2_ forms; González *et al*., 2014, 2016; Pokrovsky *et al*., 2018). Specifically, we quantified biosorption potential, buffering capacity, and functional group composition in 20 *Sphagnum* species collected from nature and transplanted on peat substrate under laboratory-controlled conditions and in 4 conspecifics *Sphagnum* species, axenically cloned in bioreactors (Beike *et al*., 2015; Heck *et al*., 2021). We then assessed gas exchange and chlorophyll fluorescence in 5 representative species (*S. auriculatum, S. balticum, S. girgensohnii, S. palustre,* and *S. strictum*), spanning major taxonomic sections. This was performed in samples under submerged conditions across a pH gradient, by using an aquatic chamber as innovative equipment, which combines the accuracy of state-of-art direct gas-exchange measurements (CO_2_ assimilation) with the sensitivity of chlorophyll fluorescence, in a highly controlled and standardized system (Hupp *et al*., 2021), overcoming the limitations of traditional approaches that rely solely on proxies such as O₂ evolution, fluorescence, or ¹⁴C incorporation (for a review, see Pedersen *et al*., 2013). Such an aquatic chamber has been recently used for algae (Hupp *et al*., 2021; Steensma *et al*., 2025) and lichens (Meyer *et al*., 2024), and, so far, in only one case for mosses (*i.e.*, *Sphagnum* spp.; Norby *et al*., 2023). However, in Norby *et al*. (2023) the moss samples were analysed immediately after field collection, without examining the effects of pH on their photosynthetic capacity or applying a standardized protocol.

Here, we optimize the analytical protocol for moss gas exchange measurements and address the following specific questions:

1. Are *Sphagnum* mosses photosynthetically efficient under waterlogging conditions, and what is the optimal pH range for that?
2. What are the differences between species in terms of photosynthetic capacity and surface chemical properties?
3. Can the chemical reactivity of *Sphagnum* surfaces be predictive of photosynthetic capacity?
4. Which species exhibit the highest potential for metal biosorption in polluted waters?
5. Are there differences between *Sphagnum* mosses collected in the field and clones grown axenically in bioreactors?

## Materials and Methods

### Moss samples

In this study, 20 *Sphagnum* species were investigated (Fig. **1**), covering the five main *Sphagnum* subgenera, *i.e., Rigida, Sphagnum, Acutifolia, Subsecunda*, and *Cuspidata.* Namely, *S. compactum* DC. and *S. strictum* Sull. from *Rigida* subg.; *S. divinum* Flatberg & Hassel, *S. centrale* C.E.O. Jensen, *S. medium* Limpr., *S. palustre* L., and *S. papillosum* Lindb. from *Sphagnum* subg.; *S. squarrosum* Crome and *S. teres* Ångstr. (*Squarrosa* section), *S. wulfianum* Girg. (*Polyclada* sect.), *S. aongstroemii* C. Hartm. (*Insulosa* sect.), *S. girgensohnii* Russow., *S. fimbriatum* Wilson, and *S. subnitens* Russow & Warnst. (*Acutifolia* sect.) from *Acutifolia* subg.; *S. auriculatum* Schimp. And *S. subsecundum* Nees. from *Subsecunda* subg.; *S. angustifolium* (Russow) C.E.O. Jensen, *S. balticum* (Russow) C.E.O. Jensen, *S. cuspidatum* Ehrh. ex Hoffm., and *S. tenellum* (Brid.) Pers. ex Brid. from *Cuspidata* subg. All species were collected in Finland, with the exception of *S. strictum* (central Sweden). Details on the collection areas, habitats, and growth form of *Sphagnum* spp. Are provided in Supporting Information Table **S1**. The *Sphagnum* species were identified through dichotomous keys in Michaelis (2019) and Laine *et al*. (2018), and by stereomicroscope and light microscopy observations.

**Fig. 1.**
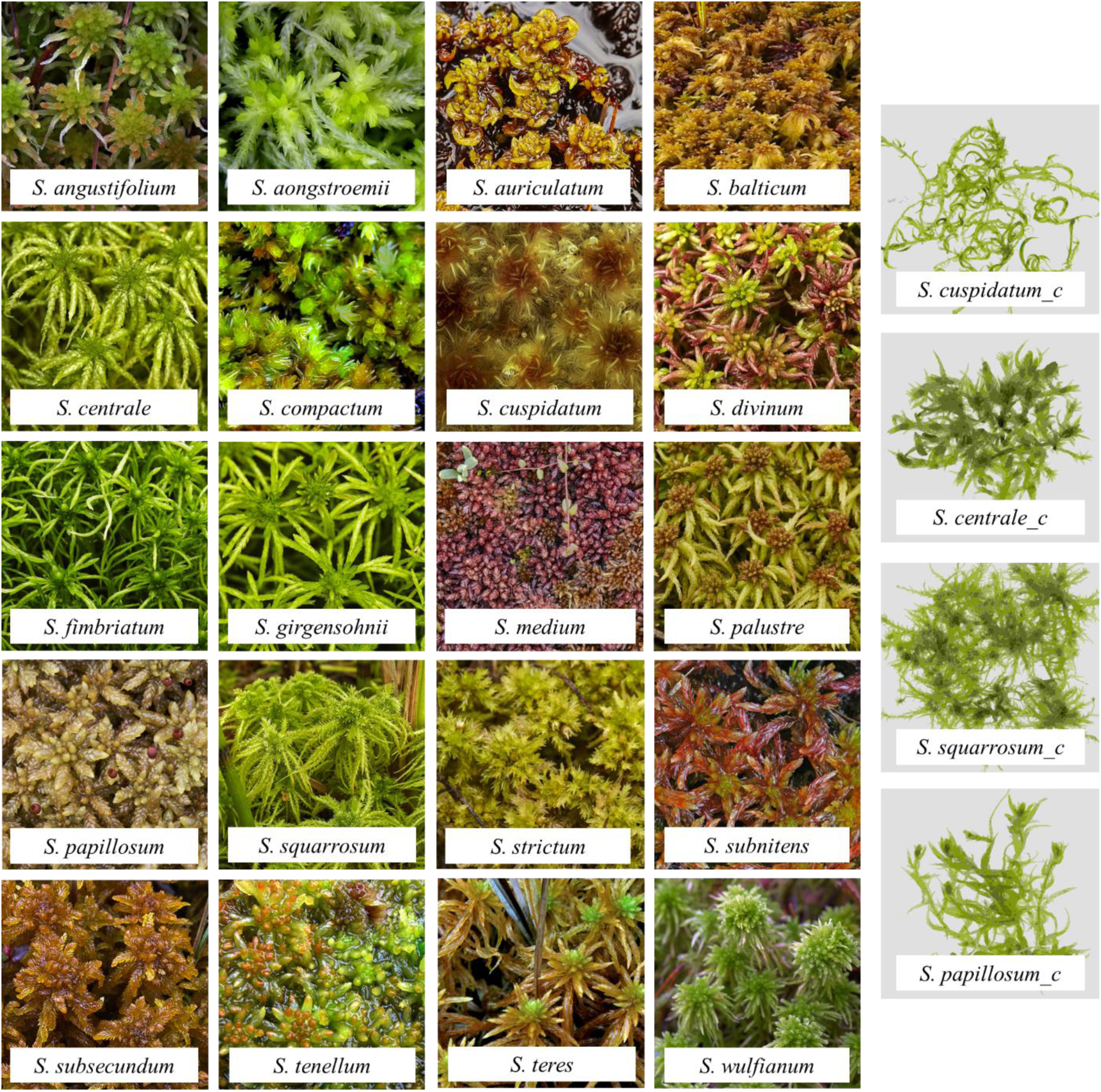
Habit photographs of the field collected and *in vitro*-cultivated (*c*) *Sphagnum* spp. used for the experiments.

After collection, the apical portions of the mosses (approximately 2 cm of stem including the capitulum) were excised from the main shoots and transplanted onto a natural substrate composed of peat soil overlaid with commercially available dead *Sphagnum* (Biomeexotic®). The dead *Sphagnum* served as a barrier to prevent direct contact between the living moss tissue and the peat soil. Cultures were maintained in a climatic chamber (Sanyo-Gallenkamp SGC970, Loughborough, UK), under controlled conditions of 20 °C, a 16/8 h light/dark photoperiod, and a photosynthetic photon flux density (PPFD) of 80 μmol m⁻² s⁻¹ to prevent tissue bleaching, etiolation, and photoinhibition in moss cultures. Moisture was maintained by covering the cultures with transparent lids and regularly misting with deionized water every three days. Additional hydration from below was ensured by placing perforated polystyrene trays containing the cultures in larger plastic containers partially filled with deionized water, allowing capillary uptake. All analyses were conducted on newly developed shoots regenerated from the transplanted gametophytes.

In addition, samples of *S. centrale*, *S. cuspidatum*, *S. papillosum*, and *S. squarrosum* cloned in sterile and axenic conditions (Heck *et al*., 2020), and available via the International Moss Stock Center IMSC (www.moss-stock-center.org), were used for comparative studies. These clones were cultivated from spores in 5 L photobioreactors (Applikon, Schiedam, The Netherlands) in liquid Knop medium (Reski & Abel, 1985) with addition of micronutrients, 2 % sucrose and ammonium nitrate, according to Beike *et al*. (2015). Photoperiod was of 16 h light to 8 h dark, with a light intensity set to 120 µmol m^-2^ s^-1^ using light tubes (Philips TLD 18 W/25) according to Hohe & Reski (2005). The pH within the bioreactors was adjusted by automatic titration with 0.5 M KOH and 0.5 M HCl, and air was insufflated with 0.3 vvm air according to Hohe & Reski (2005). Further details on cultivation method of mosses in photobioreactors are reported in Beike *et al*. (2015).

### Moss surface characterization

Moss surfaces were characterized in terms of surface charge excess and binding sites by acid–base titration at a pH range of 3-10, following González &Pokrovsky (2014). The experiments were carried out in 0.01 M NaNO_3_; the moss biomass was of 1 g_dw_ L^-1^, at room temperature (20 ± 1 °C), under an N_2_ atmosphere and continuous stirring. The pH was measured by a pH/mV meter ion analyser (Orion Lab Star PH111; Thermo Fisher Scientific, Waltham, MA, USA) with an accuracy of ± 0.005. Acid and basic titrations were conducted in triplicates for each moss, via separately adding aliquots of HCl and NaOH titrant solutions (from 0.01 to 0.1 N). Reference solutions, *i.e.,* biomass-free supernatant solution obtained after 1 h moss stirring under N_2_ bubbling, were titrated in parallel and used for calculation of charge excess values following previous methodological approaches González *et al*. (2014, 2016).

The four moss species cultivated in bioreactors were treated as the conspecific mosses grown in the climatic chamber and used for comparison purposes. Before titration, all samples were washed 3 times (10 min each washing) with filter-sterilized water (18 MW; 1 g moss/ 1 L water), as suggested in Di Palma *et al*. (2019) in order to remove excess of DOC (Dissolved Organic Carbon) released by moss that could interfere with proton and hydroxyl consumption during the titration experiment.

### CO2 exchange and chlorophyll fluorescence in waterlogged mosses

Gas exchange measurements were carried out using a LI-6800 Portable Photosynthesis Systems (LI-COR Biosciences Inc., Lincoln, NE, USA) equipped with a 6800-18 Aquatic Chamber, a novel setup which enables the simultaneous measurement of chlorophyll fluorescence and CO_2_ exchange in wet samples. Prior to our experimental measurements, we tested moss samples both suspended in water and simply moistened, with varying amount of solution added in the chamber, in order to check the performances of the aquatic chamber according to different degrees of sample moisture. Preliminary tests of light response were conducted to evaluate the response range of *Sphagnum* samples and to determine optimal measurement conditions prior to the start of the definitive experiments. Additional tests were performed to assess humidity and temperature regulation within the chamber, to ensure stable and optimal conditions during measurements.

One cultivated moss species was randomly selected for analysis from each of the five main *Sphagnum* subgenera, *i.e., S. auriculatum* (*Subsecunda* subg.)*, S. balticum* (*Cuspidata* subg.), *S. girgensohnii* (*Acutifolia* subg.), *S. palustre* (*Sphagnum* subg.), and *S. strictum* (*Rigida* subg.). For each experimental measurement, 3 to 5 green and juvenile *Sphagnum* shoots were used, each consisting of a capitulum and a 3 cm-long stem, newly generated from the mother transplants and collected after 30 to 60 days of growth, once the capitulum was fully formed. Samples exhibiting bleaching or pigmentation deviating from green were avoided, as the alteration in colour can determine lower photosynthetic rates (Maseyk *et al*., 1999).

Inside the aquatic chamber, *Sphagnum* samples were analysed while suspended in 15 mL of filter-sterilized water (18 MW), to assess their photosynthetic capacity under conditions simulating submersion. Measurements were conducted as a function of light intensity and pH, with five replicates per species. Eleven light intensities (PAR - Photosynthetically Active Radiation - ranging from 0 to 1000 μmol m^-2^ s^-1^) and four pH values (3, 5, 7, and 9) were tested. The air temperature was set to 20 °C, the air flow at 200 μmol s^-1^, and the CO_2_ concentration was maintained at 430 ppm inside the aquatic chamber. The following light intensities were used to generate A/Q response curves: 20, 40, 60, 80, 100, 200, 300, 400, 600, 800, 1000 μmol m^-2^ s^-1^. Chlorophyll fluorescence under light conditions was assessed simultaneously with gas exchange measurements to obtain values of maximum efficiency of PSII photochemistry in the light-adapted state (F_v_’/F_m_’) at each light level. In contrast, F_v_/F_m_ (maximum efficiency of PSII photochemistry in the dark-adapted state) values were obtained at the beginning of each experiment following 30 minutes of dark acclimation, according to the method described by Jassey & Signarbieux (2019). Non-photochemical quenching (NPQ) values were automatically calculated by the software embedded in the LI-6800 system. The maximum photosynthetic rate (A_max_, μmol m^−2^ s^−1^), the quantum yield of CO_2_ assimilation (*Φ*, mol mol^−1^), dark respiration (R_d_, μmol m^−2^ s^−1^), and light compensation point (LCP, μmol m^−2^ s^−1^) were estimated from individual A/PPFD response curves as reported in Tian-gen *et al*. (2017).

The pH of solution was adjusted by adding aliquots of HCl or NaOH 0.1-0.01 M, and monitored during the experiment by a pH probe inserted into the sample cuvette through the O-ring-sealed port on the side of the aquatic chamber. The pH oscillations during a measurement time of 40 min were minimal, with a maximum variation of ± 0.4 found only in one replicate.

### Data treatment

All data from titration analysis and gas exchange measurements were reported on a dry weight basis. Data analysis and statistics were performed by using Microsoft Excel-XLSTAT, Synergy Kaleidagraph v. 4.0, and MATLAB R2024b Update 6.

Normality and homogeneity of variance were assessed using the Shapiro-Wilk and Levene tests, respectively. When data were non-normally distributed, log transformation was applied prior to parametric analyses. For titration and light curve response datasets, the significance of pH and moss species/section effects was analysed via ANCOVA with Tukey test as post-hoc. The Mann-Whitney test was used to compare mosses cultivated on peat soil and cloned in bioreactors, while the Kruskall-Wallis with Dunn multiple comparison test and Bonferroni correction assessed significance of light parameters differences as a function of pH and moss species. Agglomerative hierarchical cluster analysis was applied to the acid-base titration data to explore differences among *Sphagnum* subgenera and among sections. Pearson correlations were performed to check relationships between the concentrations of surface functional groups (both total and by type) and photosynthesis parameters.

Moss surface binding site concentrations and apparent equilibrium constants from the titration experiment were estimated using the Linear Programming Model (LPM, Brassard *et al*., 1990; Smith & Ferris, 2001), following the protocol established for bacteria and mosses (González *et al*., 2016), as detailed in the Supporting Information Method **S1**.

## Results

### Moss surface characterization

Surface chemical properties of the 20 *Sphagnum* spp. were described in terms of titration curves (Fig. **2**), pH_pzc_ (values of pH corresponding to zero net proton adsorption, *i.e.,* zero charge), type and amount of surface binding sites (Table **1**, Supporting Information Fig. **S1**).

**Fig. 2.**
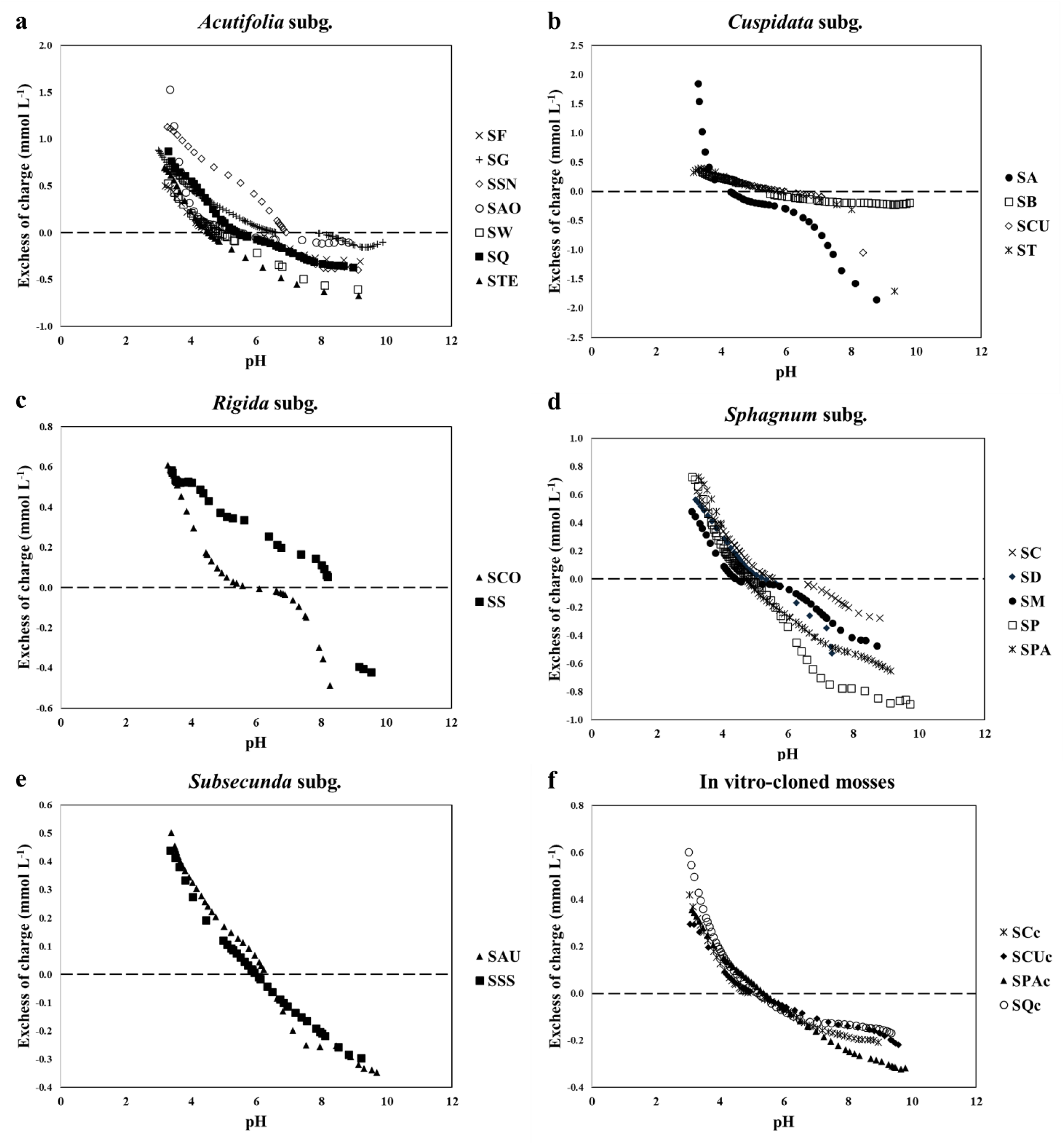
Acid-base titration curves (*n* = 3) of the 20 *Sphagnum* species collected from nature and the 4 *in vitro*-cloned mosses. SA = *S. angustifolium*; SAO = *S. aongstroemii*; SAU = *S. auriculatum*; SB = *S. balticum*; SC = *S. centrale*; SCO = *S. compactum*; SCU = *S. cuspidatum*; SD = *S. divinum*; SF = *S. fimbriatum*; SG = *S. girgensohnii*; SM = *S. medium*; SP = *S. palustre*; SPA = *S. papillosum*; SQ = *S. squarrosum*; SS = *S. strictum*; SSN = *S. subnitens*; SSS = *S. subsecundum*; ST = *S. tenellum*; STE = *S. teres*; SW = *S. wulfianum*; “c” = clone.

**Table 1.**
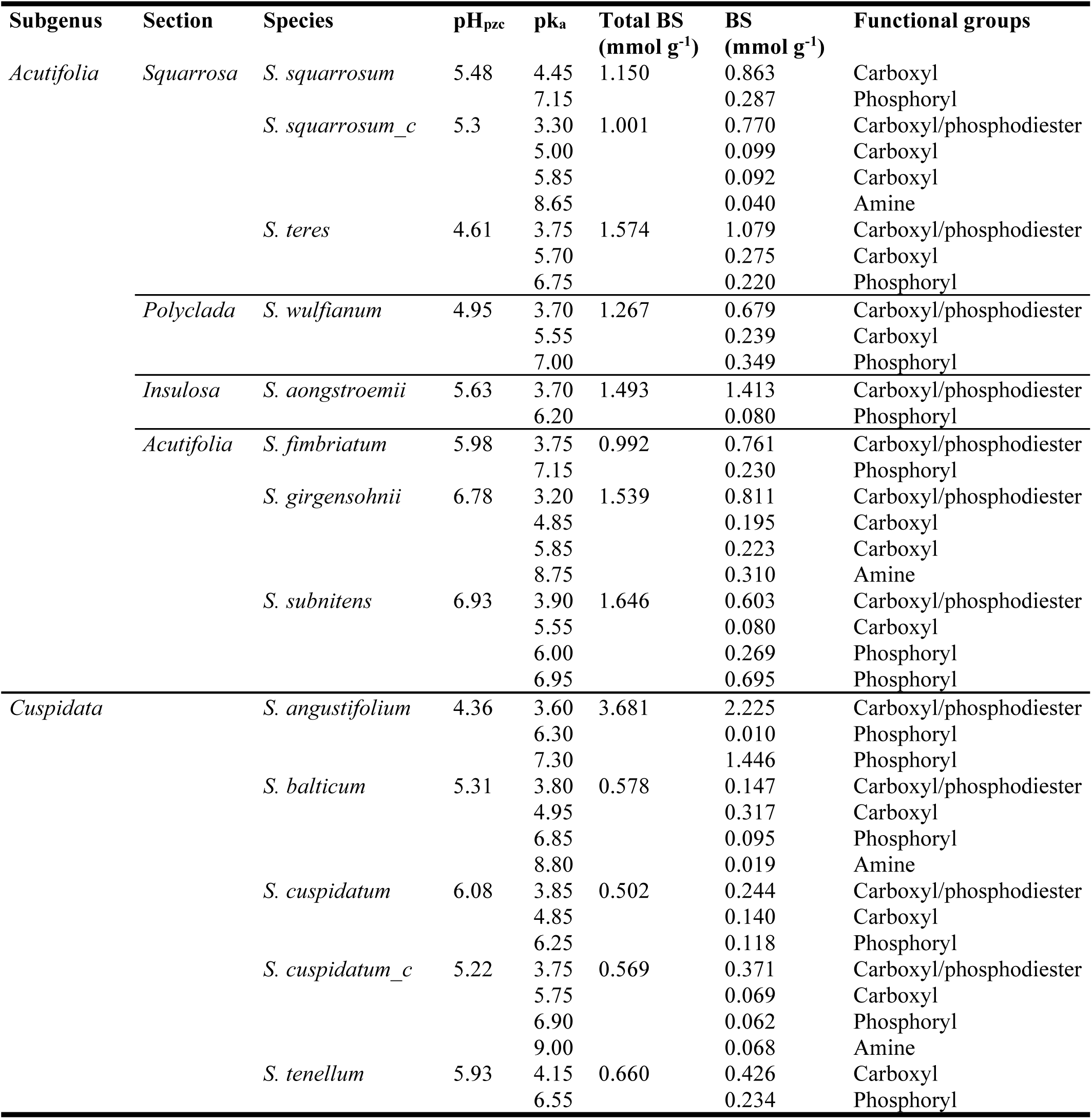
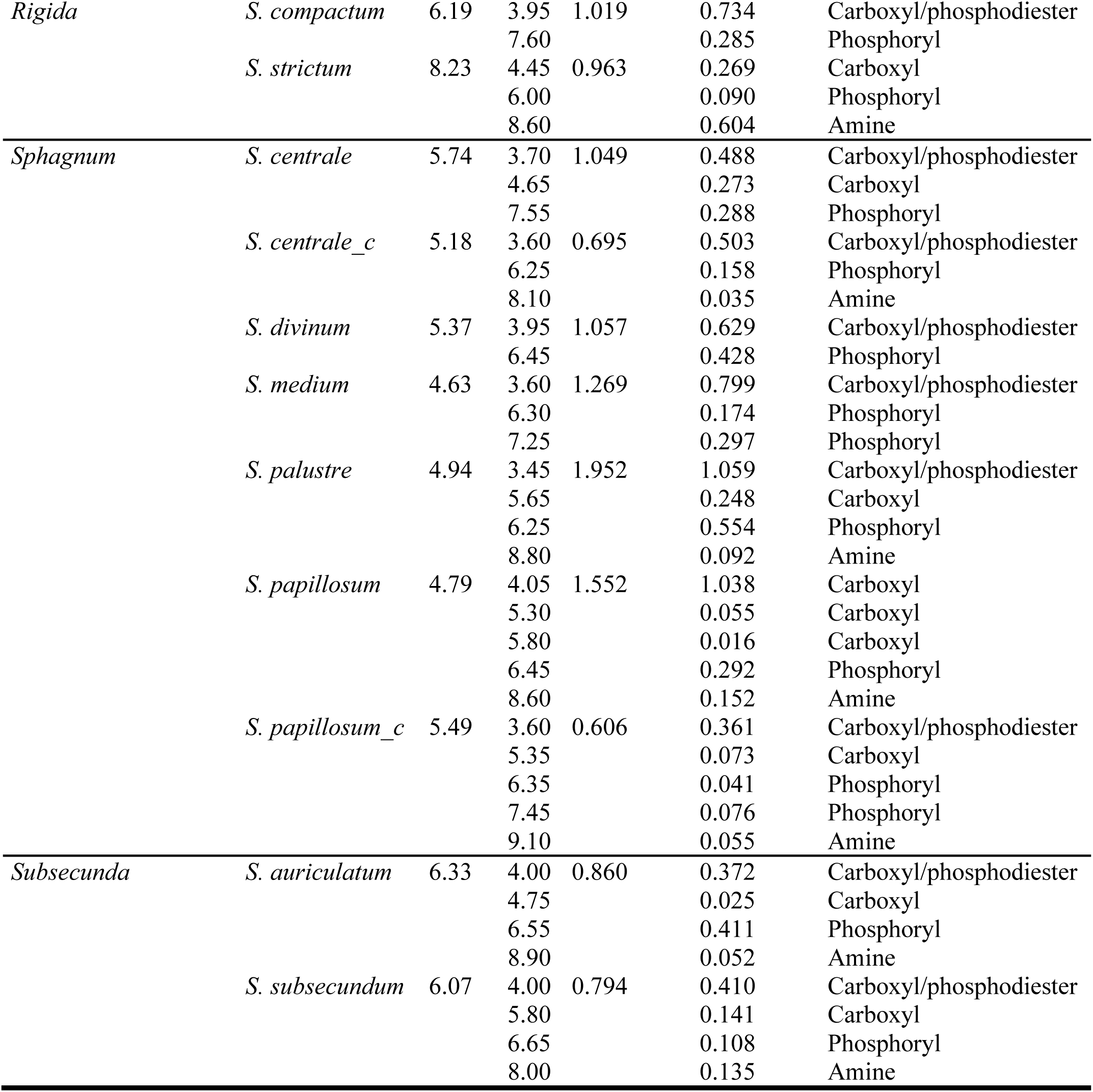
Surface acid-base titration and LPM model results for the 20 *Sphagnum* species (*n* = 3). *pH_pzc_* = pH at the point zero charges. *K_a_* = proton dissociation constant; “c” =moss cloned in photobioreactor; BS = binding sites.

Acid-base properties within the *Acutifolia* subg. differed significantly, considering *S. wulfianum* (*Polyclada* sect.) *vs S. girgenshonii* (*p* = 0.0001) and *S. subnitens* (*p* = 0.0004). For the pH range of 3.3-9.1 (Fig. **2a**), *S. subnitens* exhibited the highest proton adsorption (1.13 mmol L⁻¹), while *S. wulfianum* showed the most negative surface charge (−0.603 mmol L⁻¹), aligning with their respective pH_pzc_ values (6.93 and 4.95; Table **1**).

In *Cuspidata* subg. (Fig. **2b**), *S. angustifolium* significantly differed from *S. cuspidatum* (*p* <0.0001) and *S. tenellum* (*p* = 0.0002), with pH_pzc_ values of 4.36, 6.08 and 5.93, respectively (Table **1**). For a pH range of 3.3-8.4, the highest excess of adsorbed protons was exhibited by *S. tenellum* (0.399 mmol L^-1^), followed by *S. cuspidatum* (0.353 mmol L^-1^) and *S. angustifolium* (0.155 mmol L^-1^), whilst the highest negative surface charge was for *S. angustifolium* (−1.57 mmol L^-1^), comparable with the value for *S. cuspidatum* (−1.04 mmol L^-1^).

Within the *Sphagnum* subg. (Fig. **2d**), significant differences were found for *S. papillosum vs S. centrale* (*p* <0.0001), *S. divinum* (*p* <0.0001) and *S. palustre* (*p* = 0.0002), and for *S. palustre vs S. centrale* (*p* = 0.0068). In the pH range 3.3-8.8, the highest excess of adsorbed protons for *S. papillosum* and *S. palustre* were 0.725 and 0.568 mmol L^-1^, respectively, while the highest negative surface charge amount was −0.848 mmol L^-1^ for *S. palustre* and −0.608 mmol L^-1^ for *S. papillosum*. Although these differences, pH_pzc_ values of *S. papillosum* and *S. palustre* were similar, with values of 4.79 and 4.94, respectively (Table **1**).

The pH_pzc_ value for *S. strictum* (8.23) was 2 units higher than that of *S. compactum* (6.19; Table **1**). However, no significant differences in terms of excess of charges were found within *Rigida* subg. (Fig. **2c**), with the two moss species of this group having comparable values of highest positive and negative charge values (on average, 0.588 and −0.638 mmol L^-1^) in the pH range of 3.4-8.3. Similarly, no differences between *S. auriculatum* and *S. subsecundum* within *Subsecunda* subg. (Fig. **2e**), neither in terms of pH_pzc_ (6.33 and 6.07, respectively; Table **1**) and surface charges (on average, −0.469 and 0.69 mmol L^-1^).

Application of the LPM model revealed that all *Sphagnum* spp. had surface carboxyl/phosphodiester, carboxyl, phosphoryl, and amine groups, with a generally higher number of carboxyl/phosphodiesters and lower of amines (Table **1**; Supporting Information Fig. **S1**). By comparing all subgenera, the highest total amount of binding sites occurred in moss species of the *Acutifolia* and *Sphagnum* subg. (on average, 1.38 mmol g⁻¹), followed by *Rigida* (0.991 mmol g⁻¹), *Subsecunda* (0.827 mmol g⁻¹), and *Cuspidata* (0.435 mmol g⁻¹). Carboxyl/phosphodiesters were more abundant in *Acutifolia* and *Cuspidata* (∼ 0.88 mmol g^-1^), and in lower amount in mosses of the *Subsecunda* subg. (0.391 mmol g^-1^). The amount of carboxyl functional groups was ∼0.35 mmol g^-1^ for the *Sphagnum* and *Acutifolia* subg., ∼0.35 mmol g^-1^ for *Cuspidata* and *Rigida* subg., and 0.083 mmol g^-1^ for *Subsecunda* subg. Phosphoryls groups were in amount of ∼0.36 mmol g^-1^ for *Cuspidata*, *Acutifolia,* and *Sphagnum* subg. while for *Subsecunda* and *Rigida* subg. equal to 0.259 and 0.188 mmol g^-1^, respectively. *Rigida* subg. displayed the highest amount of amine (0.604 mmol g^-1^), while *Cuspidata* subg. the lowest (0.019 mmol g^-1^).

Within the *Acutifolia* subg., the highest number of carboxyl/phosphodiesters was displayed by *S. aongstroemii* (*Insulosa* sect.; 1.413 mmol g^-1^) and by *S. teres* (*Squarrosa* sect.; 1.079 mmol g^-1^). The other species of the same subgenus had on average 0.713 mmol g^-1^. *Sphagnum squarrosum* (*Squarrosa* sect.) did not possess carboxyl/phosphodiesters, while showing the highest amount of carboxyls (0.863 mmol g^-1^) compared to the other species of the same subgenus (0.253 mmol g^-1^, on average between *S. teres*, *S. wulfianum*, *S. girgenshonii*, *S. subnitens*). Phosphoryls, absent in *S. girgenshonii*, ranged between 0.08 (*S. aongstroemii*) and 0.964 mmol g^-1^ (*S. subnitens*), while amines were present only in *S. girgenshonii* (0.31 mmol g^-1^). Comparing sections of *Acutifolia* subg., averages of total binding sites ranged between 1.267 mmol g^-1^ (*Polyclada* sect.) and 1.493 mmol g^-1^ (*Insulosa* sect.), with comparable values for *Acutifolia* (1.392 mmol g^-1^) and *Squarrosa* (1.362 mmol g^-1^) sections. *Insulosa* sect. had the highest number of carboxyl/phosphodiesters (1.413 mmol g^-1^) followed by *Squarrosa* (1.079 mmol g^-1^). Carboxyl groups were present in *Squarrosa* sect. in averaged amount of 0.569 mmolg^-1^, and in *Acutifolia* and *Polyclada* in amount of ∼0.245 mmol g^-1^). Phosphoryl amount was highest in *Acutifolia* sect. (0.597 mmol g^-1^) and lowest in *Insulosa* sect. Amine were displayed only in *Acutifolia* sect. and only for *S. girgenshonii* (0.310 mmol g^-1^).

Within the *Cuspidata* subg., carboxyl/phosphodiester groups, absent in *S. tenellum*, were 15 and 9 times higher in *S. angustifolium* (2.225 mmol g^-1^) compared to *S. balticum* (0.147 mmol g^-1^) and *S. cuspidatum* (0.244 mmol g^-1^), respectively. Carboxylate groups were absent in *S. angustifolium* and equal to 0.294 mmol g^-1^ in the other moss species of the same subgenus (*S. balticum*, *S. cuspidatum*, *S*. *tenellum*). The highest value of phosphoryls was attributed to *S. angustifolium* (1.456 mmol g^-1^) and ranged between 0.095 (*S. balticum*) and 0.234 mmol g^-1^ (*S*. *tenellum*). Amines were found only in *S. balticum* (0.019 mmol g^-1^).

The two *Sphagnum* species of *Rigida* subg. had both phosphoryl groups (0.188 mmol g^-1^, on average). Tentative carboxyl/phosphodiester groups were found only in *S. compactum* (0.734 mmol g^-1^), while carboxylates and amines were detected only in *S. strictum*, in amounts of 0.296 and 0.604 mmol g^-1^, respectively.

Within the *Sphagnum* subg., most of possible functional groups were represented by carboxyls/phosphodiesters and phosphoryls. The first, absent in *S. papillosum*, were higher in *S. palustre* (1.059 mmol g^-1^) and equal to 0.639 mmol g^-1^, on average between *S. centrale*, *S. divinum* and *S. medium*; the latter ranged between 0.288 (*S. centrale*) and 0.554 mmol g^-1^ (*S. palustre*). Carboxyls, not present in *S. divinum* and *S. medium*, were equal to 1.109 mmol g^-1^ in SPA, an amount 4 times higher in *S. centrale* and *S. palustre* (0.26 mmol g^-1^, on average). Amines were not found in mosses of *Sphagnum* subg.

Tentative carboxylate/phosphodiester, carboxylate, phosphorylate and amine groups were found in both species of *Subsecunda* subg., in amount of 0.391, 0.083, 0.259 and 0.093 mmol g^-1^, respectively, and generally in a higher amount in *S. subsecundum* than in *S. auriculatum*, except for phosphoryls, approximately 4 times as high in *S. auriculatum* (0.411 mmol g^-1^) compared to *S. subsecundum* (0.108 mmol g^-1^).

The cluster analysis of acid-base titration data showed the overall variability in acid-base properties across subgenera, identifying six main clusters (C1–C6; Fig. **3**).

**Fig. 3.**
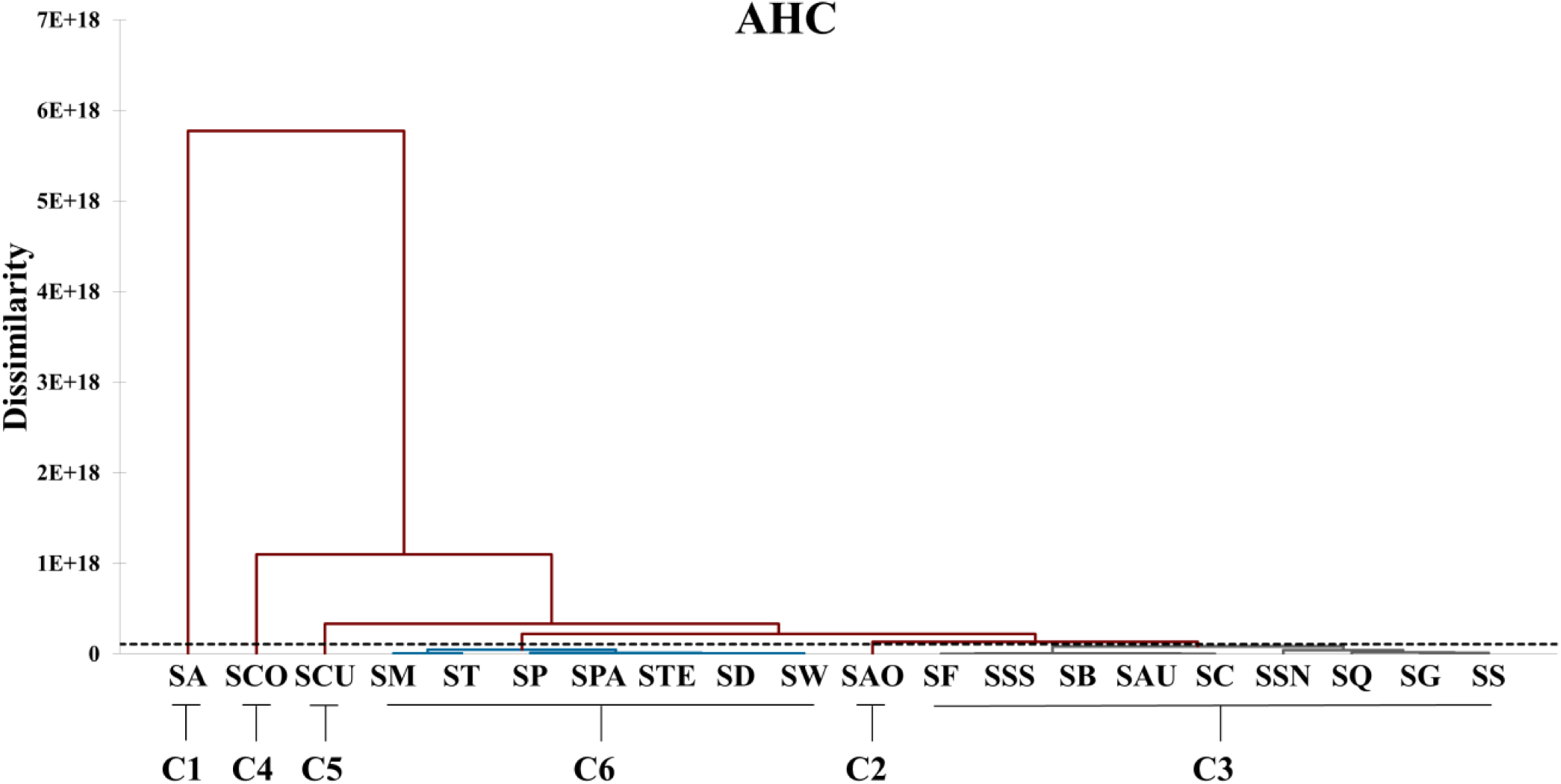
Agglomerative hierarchical cluster (AHC) analysis applied to the acid-base titration data (*n* = 3). C1-C6 = clusters; SA = *S. angustifolium*; SAO = *S. aongstroemii*; SAU = *S. auriculatum*; SB = *S. balticum*; SC = *S. centrale*; SCO = *S. compactum*; SD = *S. divinum*; SF = *S. fimbriatum*; SG = *S. girgensohnii*; SM = *S. medium*; SP = *S. palustre*; SPA = *S. papillosum*; SQ = *S. squarrosum*; SS = *S. strictum*; SSN = *S. subnitens*; SSS = *S. subsecundum*; SCU = *S. cuspidatum*; ST = *S. tenellum*; STE = *S. teres*; SW = *S. wulfianum*.

The clusters C3 and C6 broadly corresponded to subgeneric and sectional taxonomy. Namely, both *Subsecunda* species grouped in C3, while *Sphagnum* subg. mainly in C6, except *S. centrale* (C3). The *Acutifolia* subg. displayed strong clustering by section, with intra-section variation only in *Squarrosa*. For the *Cuspidata* and *Rigida* subg. significant intra-group divergence was found. *Cuspidata* species each formed separate clusters, with *S. angustifolium* forming a different group (C1), strongly distinct from the cluster C3, which includes *S. balticum* (*Cuspidata* subg.) *Subsecunda* and most *Acutifolia* spp. Similarly, *Rigida* species clustered apart, with *S. compactum* in C4 and *S. strictum* in C3.

### In vitro vs peat-grown mosses

The mosses grown in photobioreactors displayed pH_pzc_ ranging between 5.2 (*S. centrale*) and 5.5 (*S. papillosum*; Table **1**), with *S. squarrosum* clone showing the highest proton adsorption (5.49 mmol L⁻¹), and *S. papillosum* the highest negative surface charge (−3.22 mmol L⁻¹; Fig. **2f**). Compared to the clones, *S. centrale*, *S. cuspidatum*, *S. papillosum*, and *S. squarrosum* grown on peat presented the highest excess of adsorbed protons and of negative surface charge with equal pH ranges, being for most of all cases 2 times higher than those in clones (Fig. **2**). The pH_pzc_ values were similar comparing *S. centrale* and *S. squarrosum* with the corresponding clones, while for *S. papillosum* and *S. cuspidatum* there was a difference of almost one pH unit between moss cultivated on peat and the related clones (*i.e.,* 4.8 for *S. papillosum* on peat and 5.5 for cloned *S. papillosum*; 6.08 for *S. cuspidatum* and 5.22 for cloned *S. cuspidatum*; Table **1**). Despite these results, the Mann-Whitney test revealed no significant differences in terms of reactivity among clones, nor between mosses cultivated on peat with the conspecific moss grown in photobioreactors.

The total amount of tentative functional groups in clones (Table **1**) was 1.001 mmol g^-1^ for *S. squarrosum*, followed by *S. centrale* (0.695 mmol g^-1^), *S. papillosum* (0.606 mmol g^-1^) and *S. cuspidatum* (0.569 mmol g^-1^). Mosses cultivated on peat displayed a total amount of binding sites, generally higher by about 2-fold compared to conspecific clones The sole exception was for *S. cuspidatum*, as the total amount of functional groups was slightly higher in clones (0.569 mmol g^-1^) than in peat-cultivated moss (0.502 mmol g^-1^). Among clones, *S. squarrosum* had the highest number of carboxyls/phosphodiesters (0.770 mmol g^-1^) and carboxyls (0.191 mmol g^-1^), while phosphoryls and amines were found more in *S. centrale* (0.158 mmol g^-1^) and *S. cuspidatum* (0.068 mmol g^-1^), respectively (Table **1**).

Considering each specific tentative functional group (Table **1**), carboxyls/phosphodiesters were slightly higher in *S. centrale* and *S. cuspidatum* clones than in the related mosses cultivated on peat (0.503 mmol g^-1^ for cloned *S. centrale* vs 0.488 mmol g^-1^ for peat-grown *S. centrale*; 0.371 mmol g^-1^ for cloned *S. cuspidatum* vs 0.244 mmol g^-1^ for peat-grown *S. cuspidatum*). Carboxyls were absent in cloned *S. centrale* and in amount between 0.069 mmol g^-1^ (*S. cuspidatum* clone) and 0.191 mmol g^-1^ (*S. squarrosum* clone). These functional groups were higher in mosses cultivated on peat by a factor of 15, 5 and 2, respectively, for *S. papillosum*, *S. squarrosum*, and *S. cuspidatum*. Tentative phosphoryls groups, not present in cloned *S. squarrosum*, were 2-fold higher in peat-cultivated mosses (on average, 0.233 mmol g^-1^) than in conspecific clones (on average, 0.112 mmol g^-1^). Amines were present only in clones in amount ranging between 0.035 mmol g^-1^ (*S. centrale*) and 0.068 mmol g^-1^ (*S. cuspidatum*).

### Gas exchange as a function of pH

Gas exchange and chlorophyll fluorescence measurements conducted on 5 *Sphagnum* species (*S. auriculatum*, *S. balticum, S. girgensohnii*, *S. palustre*, *S. strictum*), representative of the main *Sphagnum* subgenera (*Subsecunda*, *Cuspidata*, *Sphagnum*, *Rigida*), revealed significant interspecific variability and pH-dependent differences in CO_2_ assimilation patterns. Considering the CO_2_ uptake as a function of light intensity (Fig. **4**), saturation generally occurred at PAR levels between 100 and 300 μmol m⁻² s⁻¹, depending on *Sphagnum* species and pH conditions.

**Fig. 4.**
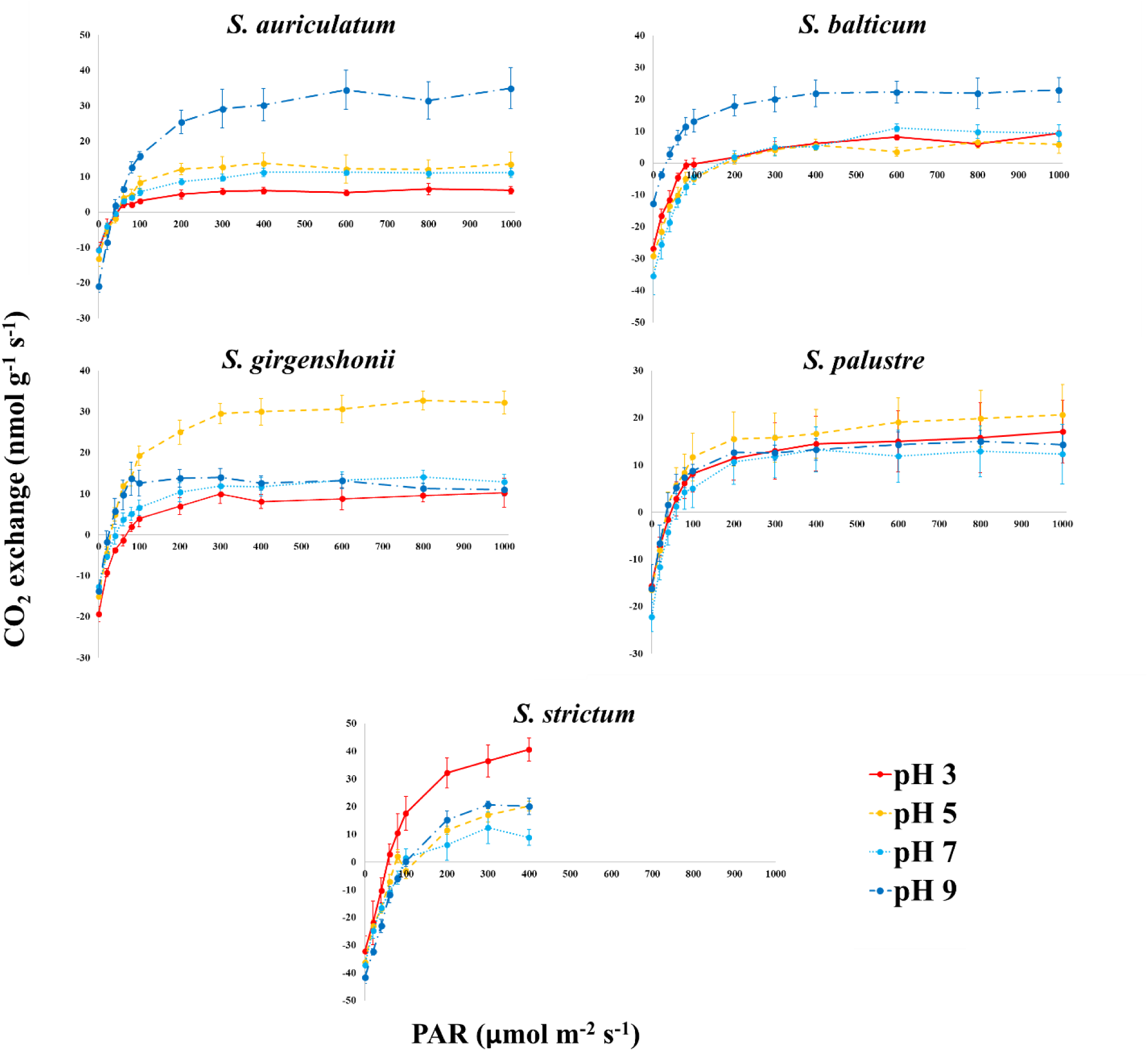
Photosynthetic light response curves (*n* = 5), shown as CO_2_ exchange rates (nmol g^-1^ s^-1^) as function of PAR (Photosynthetically Active Radiation; µmol m^-2^ s^-1^), for 5 *Sphagnum* species collected in the field and four pH values.

In *S. auriculatum* (Fig. **4**), light response curves showed a strong dependence of CO_2_ assimilation on solution pH. At low light intensities (20-40 µmol m⁻² s⁻¹ PAR), net assimilation was negative for all pH levels, with values ranging from −4.22 ± 2.20 (pH 3) to −8.53 ± 1.97 nmol g⁻^1^ s⁻¹ (pH 9). With increasing PAR, assimilation became positive and rose progressively, with a magnitude differing among treatments. The highest CO_2_ assimilation rates were exhibited by *S. auriculatum* at pH 9, reaching a value of 34.96 ± 5.78 nmol g⁻^1^ s⁻¹ at maximum irradiance (1000 µmol m⁻² s⁻¹).

By contrast, the rates of CO_2_ assimilation remained relatively low between pH 3 and 7, reaching maximum values of about 6.16, 13.8 and 11.13 nmol g⁻^1^ s⁻¹, respectively at pH 3, 5 and 7 under saturating light.

*Sphagnum balticum* exhibited patterns of light response curves comparable to those from *S. auriculatum*, with generally higher values of CO_2_ assimilation at pH 9 compared to other solution pHs (Fig. **4**). Namely, at PAR of 1000 µmol m⁻² s⁻¹, CO_2_ assimilation reached the maximum value of 22.99 ± 3.88 nmol g⁻^1^ s⁻¹ for pH 9, while it was 5.96 ± 2.85, 9.31 ± 0.83 and 9.67 ± 2.65 nmol g⁻^1^ s⁻¹, respectively for pH 3, 5 and 7. With increasing light, CO_2_ assimilation in *S. balticum* increased across all treatments with increasing PAR, but with marked differences among pH levels. At low irradiance (20-40 µmol m⁻² s⁻¹ PAR), assimilation was strongly negative under acidic and neutral conditions, with values of about −16.5 (pH 3), −21.4 (pH 5), and −25.4 (pH 7) nmol g⁻^1^ s⁻¹, and it rose more gradually for pH 3 than for pH 5 and 7. At 20 µmol m⁻² s⁻¹ PAR and pH 9, CO_2_ assimilation had a value of −3.7 ± 1.0 nmol g⁻^1^ s⁻¹, with a sharp increase up to the saturating irradiance.

In *S. girgensohnii* (Fig. **4**), the highest CO_2_ assimilation rates were detected for pH 5, with a value of about 32.22 nmol g⁻^1^ s⁻¹ at saturating light, while the lowest at pH 3 (∼10.25 nmol g⁻¹ s^-1^), followed by pH 9 (∼11.0 nmol g⁻¹ s^-1^) and pH 7 (∼12.93 nmol g⁻¹ s^-1^). At 20 µmol m⁻² s⁻¹ PAR, CO_2_ assimilation values were strongly negative at pH 3 (−9.2 ± 1.0 nmol g⁻^1^ s⁻¹) and slightly negative at pH 9 (−1.7 ± 2.7 nmol g⁻^1^ s⁻¹). Assimilation rose quickly with increasing irradiance, with the steepest increase occurring at pH 5. For equal PAR of 100 µmol m⁻² s⁻¹, CO_2_ assimilation for pH 5 reached a value of 19.27 ± 2.42 nmol g⁻^1^ s⁻¹, while it was 3.93 ± 2.04, 6.63 ± 1.90, and 12.59 ± 3.13 nmol g⁻^1^ s⁻¹ at pH 3, 7, and 9 respectively.

In *S. palustre* (Fig. **4**), CO_2_ assimilation increased with irradiance mostly uniformly among pHs, but with a different magnitude of photosynthetic activity. At 20 µmol m⁻² s⁻¹ PAR, assimilation was negative for all pH conditions, and from −11.61 ± 2.64 (pH 7) to −6.53 ± 3.83 (pH 9). At 40 µmol m⁻² s⁻¹, CO_2_ assimilation became positive for pH 5 and 9 (1.71 ± 2.59 and 1.59 ± 2.53 nmol g⁻^1^ s⁻¹, respectively), while remaining negative at pH 3 and 7. With increasing light intensity, assimilation rates rose progressively, reaching maximum values of 20.69 ± 6.42 nmol m⁻² g⁻¹ at pH 5 and 1000 µmol m⁻² s⁻¹ PAR, followed by 17.10 ± 6.66 nmol g⁻^1^ s⁻¹ at pH 3, 14.33 ± 2.27 nmol g⁻^1^ s⁻¹ at pH 9, and 12.35 ± 6.42 nmol g⁻^1^ s⁻¹ at pH 7.

For *S. strictum* (Fig. **4**), evaluation of the CO_2_ assimilation was limited up to 400 µmol m⁻² s⁻¹, as higher light intensities led to instable measurements and a progressive decline of moss vitality. At low light intensities, CO_2_ uptake remained negative across all pH levels, ranging from −19.22 ± 1.58 nmol g⁻^1^ s⁻¹ at pH 7 to −10.26 ± 4.49 nmol g⁻^1^ s⁻¹ at pH 3. The highest assimilation increase occurred at pH 3, particularly from 32.22 ± 5.44 nmol g⁻^1^ s⁻¹ at 200 µmol m⁻² s⁻¹ to 43.35 ± 4.20 nmol g⁻^1^ s⁻¹ at 400 µmol m⁻² s⁻¹. At pH 5, assimilation also increased with irradiance, with values from 11.60 ± 1.47 nmol g⁻^1^ s⁻¹ (200 µmol m⁻² s⁻¹) to 23.69 ± 2.13 nmol g⁻^1^ s⁻¹ (400 µmol m⁻² s⁻¹). Similarly, CO_2_ uptake at pH 9 increased considerably up to 29.46 ± 3.07 (300 µmol m⁻² s⁻¹) before declining at 400 µmol m⁻² s⁻¹. By contrast, at pH 7 it slightly increased with irradiance, and with values not higher than 12 nmol g⁻^1^ s⁻¹.

### Light parameters

To better characterize the photosynthetic performance of the 5 moss species under different pH conditions, photosynthetic parameters were calculated from light-response curves, *i.e.,* the maximum photosynthetic rate (A_max_), the apparent quantum yield (*Φ*), the light compensation point (LCP), and the dark respiration rate (R_d_; Table **2**).

**Table 2.** Photosynthesis parameters calculated from light response curves and dark-adapted maximum quantum yield of PSII (F_v_/F_m_) for 5 *Sphagnum* species collected in the field and four pHs. Values (*n* = 5) are given with their standard errors. A_max_ =maximum photosynthetic rate; *Φ* = apparent quantum yield; LCP = light compensation point; R_d_ = dark respiration rate. Letters a and b indicate statistical significance (*p* =0.0083; Kruskal Wallis test + Dunn multiple comparison and Bonferroni correction).

| Species | pH | $A_{\max}$<br>nmol g<sup>-1</sup> s<sup>-1</sup> | $\Phi$ (x 10<sup>-4</sup>)<br>mol mol<sup>-1</sup> | LCP<br>µmol mol<sup>-1</sup> | $R_d$<br>µmol g<sup>-1</sup> s<sup>-1</sup> | $F_v/F_m$ |
| --- | --- | --- | --- | --- | --- | --- |
| *S. auriculatum* | 3 | 8.80 ± 2.86<sup>b</sup> | 1.41 ± 0.36 | 47.36 ± 6.89 | -6.60 ± 2.05 | 0.720 ± 0.028 |
|  | 5 | 14.10 ± 3.11<sup>ab</sup> | 2.60 ± 0.93 | 48.93 ± 8.03 | -11.38 ± 3.74 | 0.683 ± 0.009 |
|  | 7 | 10.91 ± 0.87<sup>ab</sup> | 1.17 ± 0.21 | 57.36 ± 6.87 | -6.82 ± 1.58 | 0.709 ± 0.005 |
|  | 9 | 30.42 ± 5.35<sup>a</sup> | 2.80 ± 0.37 | 41.02 ± 3.94 | -11.54 ± 2.21 | 0.694 ± 0.031 |
| *S. balticum* | 3 | 7.56 ± 1.46<sup>ab</sup> | 2.40 ± 0.24 | 85.07 ± 4.78<sup>ab</sup> | -20.40 ± 2.27<sup>ab</sup> | 0.643 ± 0.037 |
|  | 5 | 5.45 ± 1.59<sup>b</sup> | 2.00 ± 0.32 | 124.77 ± 18.41<sup>b</sup> | -23.06 ± 2.20<sup>ab</sup> | 0.587 ± 0.033 |
|  | 7 | 6.82 ± 2.31<sup>ab</sup> | 2.60 ± 0.68 | 123.56 ± 20.27<sup>ab</sup> | -28.08 ± 4.47<sup>b</sup> | 0.620 ± 0.025 |
|  | 9 | 19.62 ± 4.47<sup>a</sup> | 2.20 ± 0.37 | 49.97 ± 15.37<sup>a</sup> | -10.00 ± 3.00<sup>a</sup> | 0.699 ± 0.015 |
| *S. girgensohnii* | 3 | 7.83 ± 2.14<sup>b</sup> | 1.44 ± 0.34<sup>ab</sup> | 111.27 ± 38.63<sup>b</sup> | -11.26 ± 0.88 | 0.675<sup>b</sup> ± 0.025 |
|  | 5 | 28.50 ± 3.99<sup>a</sup> | 2.80 ± 0.37<sup>ab</sup> | 34.13 ± 6.03<sup>ab</sup> | -9.40 ± 1.79 | 0.697<sup>ab</sup> ± 0.017 |
|  | 7 | 8.84 ± 3.63<sup>ab</sup> | 1.02 ± 0.43<sup>b</sup> | 45.62 ± 9.63<sup>ab</sup> | -4.22 ± 1.66 | 0.720<sup>ab</sup> ± 0.008 |
|  | 9 | 19.80 ± 5.11<sup>ab</sup> | 3.00 ± 0.32<sup>a</sup> | 24.6 ± 8.69<sup>a</sup> | -6.92 ± 2.11 | 0.828<sup>a</sup> ± 0.055 |
| *S. palustre* | 3 | 23.62 ± 9.80 | 2.60 ± 0.68 | 49.94 ± 18.91 | -9.52 ± 1.84 | 0.753 ± 0.009 |
|  | 5 | 25.04 ± 8.78 | 2.60 ± 0.52 | 46.28 ± 10.54 | -10.12 ± 1.37 | 0.732 ± 0.038 |
|  | 7 | 22.76 ± 10.38 | 2.80 ± 0.66 | 65.65 ± 19.66 | -13.64 ± 2.00 | 0.768 ± 0.006 |
|  | 9 | 18.88 ± 4.55 | 2.20 ± 0.37 | 39.70 ± 8.09 | -9.20 ± 3.09 | 0.774 ± 0.005 |
| *S. strictum* | 3 | 40.65 ± 4.15<sup>a</sup> | 5.00 ± 0.95 | 61.33 ± 10.42 | -30.94 ± 7.22 | 0.582<sup>ab</sup> ± 0.025 |
|  | 5 | 20.31 ± 1.76<sup>ab</sup> | 4.80 ± 0.37 | 71.90 ± 3.39 | -34.62 ± 3.33 | 0.502<sup>b</sup> ± 0.010 |
|  | 7 | 12.40 ± 2.84<sup>b</sup> | 3.60 ± 0.68 | 100.08 ± 16.51 | -31.66 ± 1.31 | 0.611<sup>a</sup> ± 0.016 |
|  | 9 | 20.15 ± 2.93<sup>ab</sup> | 4.40 ± 0.24 | 90.46 ± 4.12 | -39.50 ± 1.35 | 0.588<sup>ab</sup> ± 0.021 |

Maximum net CO_2_ assimilation (Aₘₐₓ) varied markedly among species and pH levels (Table **2**). The highest A_max_ was observed in *S. strictum* at pH 3 (40.65 ± 4.15 nmol g⁻^1^ s⁻¹) and in *S. auriculatum* at pH 9 (30.42 ± 5.35 nmol g⁻^1^ s⁻¹), followed by *S. girgensohnii* (28.50 ± 3.99 nmol g⁻^1^ s⁻¹ at pH 5), *S. palustre* (25.04 ± 8.78 nmol g⁻^1^ s⁻¹ at pH 5) and *S. balticum* (19.62 ± 0.004 nmol g⁻^1^ s⁻¹ at pH 9). In *S. auriculatum*, A_max_ increased significantly with pH, ranging from 8.80 ± 2.86 nmol g⁻^1^ s⁻¹ at pH 3 to a maximum of 30.42 ± 5.35 nmol g⁻^1^ s⁻¹ at pH 9 (*p* < 0.0083). Similarly, *S. balticum* exhibited the highest Aₘₐₓ at pH 9 (19.62 ± 4.47 nmol g⁻^1^ s⁻¹), and almost low Aₘₐₓ values at other pHs (5.45 - 7.56 nmol g⁻^1^ s⁻¹). In *S. girgensohnii*, no clear trend was observed for A_max_ values as function of pH, with highest A_max_ recorded at pH 5 (28.50 ± 3.99 nmol g⁻^1^ s⁻¹), significantly higher than at pH 3 (7.83 ± 2.14 nmol g⁻^1^ s⁻¹; *p* < 0.0083). By contrast, *S. palustre* showed no significant differences in Aₘₐₓ across pH treatments, with values ranging from 18.88 ± 4.55 to 25.04 ± 8.78 nmol g⁻^1^ s⁻¹. In *S. strictum*, Aₘₐₓ reached the highest value at pH 3 (40.65 ± 4.15 nmol g⁻^1^ s⁻¹), significantly higher than at pH 7 (12.40 ± 2.84 nmol g⁻^1^ s⁻¹; *p* < 0.0083).

The apparent quantum yield (*Φ*) followed species-specific patterns and varied across pH treatments (Table **2**). *Sphagnum auriculatum* showed a fluctuating trend with a minimum at pH 7 (1.17 ± 0.21 × 10⁻⁴) and a maximum at pH 9 (2.80 ± 0.37 × 10⁻⁴). However, no statistically significant differences were detected among pH levels. In *S. balticum*, *Φ* exhibited relatively stable yields across different pH levels, ranging between 2.0 and 2.6 × 10⁻⁴, and no significant differences were observed. *Sphagnum girgensohnii* showed its highest *Φ* at pH 9 (3.00 ± 0.32 × 10⁻⁴), whereas the value dropped to 1.02 ± 0.43 × 10⁻⁴ at pH 7 (*p* < 0.0083). For *S. palustre*, *Φ* values were consistently high, between 2.2 and 2.8 × 10⁻⁴, and did not vary significantly with pH. *Sphagnum strictum* clearly differed from the other four moss species, showing the highest *Φ* values overall (3.6 - 5.0 × 10⁻⁴), slightly higher under acidic conditions, although no significant differences were observed among pH treatments.

The light compensation point (LCP) showed a significant pH-dependent variability in case of *S. balticum* and *S. girgenshonii* (Table **2**). In particular, *S. balticum* showed the highest compensation points overall, with extremely elevated values at pH 5 (124.8 ± 18.4 µmol mol⁻¹), while decreasing at pH 9 (49.9 ± 15.4 µmol mol⁻¹; *p* < 0.0083). Similarly, S. *girgensohnii* had LCP values declining markedly from 111.27 ± 38.63 µmol mol⁻¹ at pH 3 to 24.60 ± 8.69 µmol mol⁻¹ at pH 9 (*p* < 0.0083). For *S. auriculatum, S. palustre* and *S. strictum*, the LCP values remained relatively constant across treatments. In *S. auriculatum*, they ranged from 41.0 ± 3.9 (pH 9) to 57.4 ± 6.9 µmol mol⁻¹ at (pH 7), similarly to those observed for *S. palustre* (39.7 - 65.6 µmol mol⁻¹). Instead, *S. strictum* showed consistently high LCPs, ranging from 61.3 ± 10.4 (pH 3) to 100.1 ± 16.5 (pH 7).

Differences were observed among species in terms of respiration rates (R_d_), but with no statistical significance as function of pHs, except for *S. balticum*. In this species, higher respiration was exhibited from acidic to neutral conditions (–20.40, −23.06, −28.08 µmol g⁻^1^ s⁻¹ at pH 3, 5 and 7, respectively), with a decreasing rate at pH 9 (−10.00 µmol g⁻^1^ s⁻¹). *Sphagnum auriculatum*, *S. girgenshonii*, and *S. palustre* had the lowest respiration rates, with R_d_ values ranging from −4.22 µmol g⁻^1^ s⁻¹ (pH 7; *S. girgensohnii*) to −13.64 µmol g⁻^1^ s⁻¹ (pH 7; *S. palustre*). Instead, *S. strictum* had the highest respiration rates among species, ranging from −30.94 ± 7.22 µmol g⁻^1^ s⁻¹ at pH 3 to −39.50 ± 1.35 µmol g⁻^1^ s⁻¹ at pH 9.

The maximum quantum efficiency of PSII in dark-adapted samples (F_v_/F_m_) showed moderate variation among *Sphagnum* species and across pH treatments (Table **2**). Values ranged between 0.50 and 0.83, with most species exhibiting F_v_/F_m_ above 0.70, with a tendency to be lower under acidic conditions (pH 3–5) and higher at neutral to slightly alkaline pH (7–9). This trend was particularly evident in *S. girgensohnii*, which showed a significant (*p* < 0.0083) and progressive increase in F_v_/F_m_ from 0.675 ± 0.025 at pH 3 to 0.828 ± 0.055 at pH 9, the highest value among all species. In contrast, *S. strictum* displayed the lowest overall F_v_/F_m_ values, ranging between 0.50 (pH 5) and 0.61 (pH 7; *p* < 0.0083). The F_v_/F_m_ values for *S. auriculatum*, *S. balticum*, and *S. palustre* were maintained relatively stable across treatments, higher in *S. palustre* (0.732 - 0.774), followed by *S. auriculatum* (0.683 - 0.720) and *S. balticum* (0.587 - 0.699).

Pearson correlation analyses revealed distinct, pH-dependent relationships between the amount of surface binding sites and photosynthesis parameters (Supporting Information Table **S2**). Total binding capacity showed generally weak correlation with all traits, generally explaining less variance than individual functional groups. Considering A_max_, the strongest positive correlations were observed with carboxyl-phosphodiesters (*r* = 0.73–0.78, *r²* = 0.54–0.61) and phosphoryl groups (*r* = 0.57–0.67, *r²* = 0.33–0.44) under acidic to neutral conditions (pH 5–7). Amines were positively correlated only at pH 3 (*r* = 0.76, *r²* = 0.57), while carboxyls were generally weakly or negatively correlated, becoming strongly negatively correlated to A_max_ at pH 9 (*r* = –0.73, *r²* = 0.53). For quantum yield (*Φ*), strong correlations resulted with carboxyl-phosphodiesters at pH 5 (*r* = 0.73, *r²* = 0.5) and with amines across all pH conditions (*r* = 0.74–0.94), with a variance up to 90% (*r²*= 0.5–0.9 at pH 5 and 9). Phosphoryl groups displayed strong negative correlations with LCP across all pH levels (*r* = –0.71 to –0.88, *r²* = 0.5–0.8). Carboxyls correlated positively to LCP at pH 3 (*r* = 0.83, *r²* = 0.7), while amines positively correlated only under alkaline conditions (pH 9, *r* = 0.66, *r²* = 0.4). Strong positive correlations were found between R_d_ and phosphoryls at pH 3–7 (*r* = 0.85–0.89, *r²* = 0.7–0.8), moderate with carboxyl-phosphodiesters (*r* = 0.52–0.78, *r²* = 0.3–0.6) and negative with amines, particularly at pH 9 (*r* = –0.83, *r²* = 0.7).

Fluorescence measurements were performed in parallel with the light-response curve processing, and expressed in terms of non-photochemical quenching (NPQ) and maximum efficiency of PSII photochemistry in the light-adapted state (F_v_’/F_m_’; Supporting Information Fig. **S2**). For all examined *Sphagnum* species, NPQ rose with increasing light intensity, and followed a different pattern as function of pH (*p* < 0.0001; ANCOVA + Tukey test), except for *S. palustre*, in which NPQ activation appeared unaffected by pH changes. The NPQ curves for *S. auriculatum*, *S. balticum*, and *S. girgensohnii* showed significant differences at pH 9 compared to other pH levels (*p* < 0.0001). *Sphagnum auriculatum* showed generally higher NPQ values for intermediate pHs, exceeding 2 at 1000 µmol m^-2^ s^-1^. In *S. balticum*, the NPQ curves recorded at pH 5 and 7 differed significantly (*p* = 0.0022), particularly at high light intensities (>400 µmol m⁻² s⁻¹), reaching the highest values (2.8 ± 0.2) at pH 9 and maximum tested intensity. A similar trend was observed in *S. girgensohnii*, showing the highest NPQ response at pH 9, even if less pronounced than in *S. balticum*. In *S. strictum*, NPQ values were generally in lower ranges than in the other four examined *Sphagnum* species, < 0.3 up to 100 µmol m⁻² s⁻¹ then progressively increasing, with significant different patterns between pH 9 and pH 5 - 7, and between pH 7 and pH 3 – 5 (p < 0.0001).

In all species and for all pHs, the F_v_’/F_m_’ ratios decreased with increasing light intensity, with a steeper decline occurring within the first 200 µmol m^-2^ s^-1^ PAR. This trend did not differ as function of pH variation in most of cases, while it appeared strongly influenced by pH conditions in case of *S. strictum* (p < 0.0001; ANCOVA + Tukey test).

## Discussion

### Moss surface chemical reactivity

Titration experiments showed that peat-grown and *in vitro*-cloned *Sphagnum* spp. have strongly amphoteric surfaces controlled by dissociation of organic functional groups, identified by LPM analysis as carboxyl/phosphodiesters, carboxyls, phosphoryls, and amines. These groups derive from cell-wall components such as cellulose, humic and fulvic acids, and aliphatic and phenolic compounds (including lignin-like polymers; Renault *et al*., 2017; Bengtsson *et al*., 2018; Hodgkins *et al*., 2018; Astolfi *et al*., 2024). Their dissociation enables *Sphagnum* to modulate its local microenvironment by acidification, and influences metal adsorption and carbon dynamics (Clymo, 1973; Fein *et al*., 2001; Borrok & Fein, 2004; Gélabert *et al*., 2004). Because *Sphagnum* lacks a well-developed conducting system (Brown, 1982; Reski, 1998) and has porous leaves with hyaline cells, both external and internal cell surfaces of the entire gametophyte likely contribute to ion exchange through rapid adsorption followed by slower diffusion-controlled uptake (Qin *et al*., 2006).

Across all taxa, carboxyl/phosphodiester and carboxyl groups dominate proton- and metal- binding (Ringqvist *et al*., 2002; Ringqvist & Öborn, 2002; Pokrovsky *et al*., 2005; Vijayaraghavan & Yun, 2008; González & Pokrovsky, 2014; Di Palma *et al*., 2019; Pokrovsky *et al*., 2024). Phosphoryls likely participate in Ca²⁺ and Mg²⁺ binding and buffering under circum-neutral pH, supporting ion selectivity in minerotrophic conditions, while amines, detected in specific taxa (*S. auriculatum, S. balticum*, *S. girgensohnii*, *S. palustre*, *S. papillosum*, *S. strictum*, *S. subsecundum*), may aid cation exchange or ammonium uptake under particular environmental constraints (*e.g.,* extreme pH conditions and low metal concentrations; Mishra *et al*., 2010).

In the *Acutifolia* subg. functional-group profile reflected sectional taxonomy: *Insulosa* and *Squarrosa* species were enriched in carboxyl/phosphodiesters, whereas the *Acutifolia* section had more phosphoryls, and amines in *S. girgenshonii*. These findings suggest that differences in chemical composition of cell walls can occur even at the sectional level. Within this subgenus, the pronounced variability in acid–base behaviour and surface functional group composition occurring among species lead to important implications for their chemical reactivity, ecological distribution and contributions to peatland biogeochemistry. Species such as *S. subnitens*, with the highest pH_pzc_ and proton adsorption capacity, have a high density of deprotonatable groups and strong buffering potential under alkaline conditions, an advantage in minerotrophic habitats typically occupied by *Acutifolia* species (Supporting Information Table **S1**), which allows them to maintain low external pH even under elevated concentrations of base cations. In contrast, *S. wulfianum*, with the lowest pH_pzc_ and the most negative surface charge, can display a high affinity for cations (Ca²⁺, Mg²⁺, NH₄⁺, Na⁺, trace metals), and a potentially good adaptability to ombrotrophic, nutrient-poor environments, despite its common occurrence in minerotrophic sites (Supporting Information Table **S1**). Experiments by Granath *et al*. (2010) showed that bog species tolerate minerotrophic waters more than expected, while Johnson *et al*. (2014) demonstrated that ionic niche preferences in *Sphagnum* are evolutionarily labile and do not consistently follow phylogenetic relationships, *i.e.*, while individuals of one species occupy similar microhabitats, related species differ markedly in tolerances, and distantly related species may converge on similar niches. This suggests that microhabitat preference is shaped more by phenotypic plasticity and ecological interactions than by deep phylogenetic constraints.

In the *Cuspidata* subg., the overall total binding site density was relatively low (0.435 mmol g⁻¹), but species differed markedly. *Sphagnum angustifolium* emerged as the most chemically reactive taxon, showing the highest chemical reactivity, with the strongest negative surface charge (–1.57 mmol L⁻¹) and an exceptionally high abundance of carboxyl/phosphodiesters and phosphoryls. This profile indicates a strong capacity for binding cations, especially at neutral to slightly acidic pH. Its low pH_pzc_ (4.36) further suggests that surfaces remain negatively charged across a wide pH range, enhancing electrostatic interactions. Consequently, *S. angustifolium* represents a promising biosorbent under neutral to slightly acidic conditions, and a strong competitor in ion-rich or weakly minerotrophic habitats, consistent with its typical occurrence in oligo- to intermediate minerotrophic environments (Supporting Information Table **S1**). By contrast, *S. cuspidatum* and *S. tenellum* had higher pH_pzc_ values and lower site densities, indicating weaker cation-exchange and buffering capacities, matching their preference for very acidic, nutrient-poor hollows.

The *Sphagnum* subg. displayed one of the highest overall site densities (1.38 mmol g⁻¹), dominated by carboxyl/phosphodiester and phosphoryl groups. *Sphagnum palustre* combined a high carboxyl/phosphodiester content (1.059 mmol g⁻¹) and substantial carboxyl concentration (1.109 mmol g⁻¹) with a consistently negative surface charge (–0.848 mmol L⁻¹), supporting its strong proton adsorption and metal-binding capacity. Although often described as mesotrophic (Supporting Information Table **S1**), its chemistry suggests strong performance in ombrotrophic or acidic minerotrophic settings where proton release and cation retention confer competitive advantages. By contrast, *S. papillosum* lacked carboxyl/phosphodiester groups and consequently exhibited more limited binding versatility despite similar charge magnitudes. As a consequence, *S. papillosum*, despite its relatively high proton adsorption (0.568 mmol L⁻¹), may be less effective in acidification conditions, and more favoured in minerotrophic sites, characterized by non-extreme pHs, moderate ion-binding demand and low selective pressure to develop higher density of cation exchange sites (Vitt, 1994; Rydin & Jeglum, 2013). The absence of amines across the subgenus limits anion-binding capacity, suggesting that the chemical interactions in *Sphagnum* subg. mosses are prevalently based on cation exchange.

In contrast to the other studied subgenera, *Rigida* and *Subsecunda* subg. were more uniform in terms of chemical reactivity. Within *Rigida*, *S. strictum* and *S. compactum* showed similar overall charge behaviour, though their functional group distributions differed. The higher pH_pzc_ (8.23) of the former suggests reduced ability in proton release and weaker cation binding under acidic conditions compared to *S. compactum,* but also indicates greater suitability for alkaline, minerotrophic habitats, consistent with its common fen occurrence (Supporting Information Table **S1**). Both species possessed phosphoryl groups, but carboxylates and amines occurred only in *S. strictum*, while carboxyl/phosphodiesters were exclusive to *S. compactum*. These differences imply that *S. compactum* may bind multivalent cations more efficiently, whereas the elevated amine content in *S. strictum* confers a more amphoteric surface and potentially broader metal-binding capacity under variable pH. *Subsecunda* species exhibited balanced functional-group compositions, moderate pH_pzc_ values (6.07–6.33), and relatively weak proton adsorption, consistent with their occurrence in transitional oligo- to mesotrophic habitats (Supporting Information Table **S1**). Overall, *S. subsecundum* showed higher total site abundances than *S. auriculatum*, except for phosphoryl groups, which were ∼4-fold more abundant in *S. auriculatum*. This suggests that *S. subsecundum* possesses a broader cation-exchange capacity, whereas *S. auriculatum* may specialize in binding multivalent cations via phosphoryls. Despite these compositional differences, both species had similar pH_pzc_ and surface charges, maintaining negatively charged cell walls across typical peatland pH ranges and enabling consistent cation retention and moderate acidification. These traits likely contribute to niche differentiation within *Subsecunda*, with *S. subsecundum* better suited to slightly nutrient-poor minerotrophic conditions (Supporting Information Table **S1**) and *S. auriculatum* to environments where interactions with multivalent cations are more prominent. Overall, both species appear adapted to moderately acidic, base-influenced peatlands through similar charge behaviour but distinct functional-group strategies.

The diversity of functional groups within *Acutifolia* and *Sphagnum* explains their overall higher biosorption potential compared to *Cuspidata*, *Rigida*, and *Subsecunda*. Our results suggest that *S. angustifolium, S. subnitens* and *S. palustre* are the most efficient adsorbents of metal pollutants among the 20 *Sphagnum* species studied here. Furthermore, the exceptional performance of *S. angustifolium* (*Cuspidata*) illustrates that intra-subgeneric variability can exceed inter-subgeneric trends, underlining the importance of species-level screening for applied biosorption purposes.

The overall marked differences among *Sphagnum* species and subgenera in pH_pzc_, surface charge, and functional-group composition likely reflect underlying biochemical variation (Limpens *et al*., 2017; Bengtsson *et al*., 2018). These chemical divergences contribute to the ecological versatility of the genus and help explain species zonation, competitiveness, and roles in peatland biogeochemistry (Clymo, 1973; Rydin et al., 2006; Rydin & Jeglum, 2013). Species with higher densities of H⁺/cation-exchange groups, such as carboxyls and phenolics, can more effectively acidify their surroundings and retain nutrients, enhancing competitive performance (Glime, 2021). Cluster analysis (Fig. **3**) further showed pronounced dissimilarities even among closely related taxa (*e.g.,* within *Cuspidata* and *Rigida*), indicating species-specific surface reactivity driven primarily by cell-wall chemistry rather than anatomy. Conversely, similarities within *Subsecunda*, *Sphagnum*, and at the sectional level in *Acutifolia* suggest subgenus-level chemical traits that are conserved from early developmental stages and maintained even in artificial cultivation. This aligns with Limpens *et al*. (2017), who reported phylogenetic conservation in carbohydrate composition, while elemental and non-carbohydrate fractions are more influenced by environmental and microtopographic variation.

### In vitro-cloned mosses

*In vitro-*cloned mosses showed pH_pzc_ values similar to field-grown *Sphagnum*, confirming that laboratory cultivation preserves fundamental surface characteristics. Among clones, *S. squarrosum* displayed the highest proton adsorption capacity (5.49 mmol L⁻¹), while *S. papillosum* exhibited the highest negative surface charge (–3.22 mmol L⁻¹), suggesting species-specific differences in cell wall reactivity even under controlled cultivation conditions.

Compared to *in-vitro* clones, peat-grown mosses generally had higher functional group abundance and proton adsorption, reflecting enhanced ion-exchange capacity and buffering under natural conditions. Environmental factors such as nutrient availability, microbial interactions, and substrate chemistry, along with the developmental origin of tissues, likely influence cell wall composition. Clones, originating from spores, exhibit *de novo* tissue formation with less structural complexity and fewer or no microbial associations (Kostka *et al*., 2016; Parsons *et al*., 2025), and, consequently, may express a more limited range of cell wall polymers and functional moieties, reflecting early developmental stages and adaptation to nutrient-controlled, sterile conditions. Conversely, peat-grown shoots develop from existing gametophytic tissue and thus may retain mature morphological and biochemical features, including more developed polysaccharide composition and higher surface reactivity.

With few exceptions (*i.e.*, the absence of phosphoryl groups in cloned *S. squarrosum* and the presence of amino groups only in clones), photobioreactor-grown mosses maintained the baseline functional groups of peat-grown conspecifics. This suggests that *in vitro* cultivation preserves the fundamental surface chemistry of *Sphagnum* tissues. However, chemical differentiation among species was generally reduced in clones, likely because each clone derives from a single spore and is grown under uniform, sterile conditions (Beike *et al*., 2015; Di Palma *et al*., 2017), minimizing the environmental drivers of variation present in field-grown mosses. Lower surface reactivity in clones may also result from the absence of natural inputs, such as organic matter, aerosols, or microbial activity, that may promote the development of reactive sites and influence surface oxidation states. The exception observed for *S. cuspidatum*, where clone values slightly exceeded those of peat-grown samples, may reflect species-specific responses to substrate composition or growth environment. Normalizing site numbers to specific surface area (SSA_BET_), which is roughly twice as high in *in-vitro* clones, may partly account for differences in surface reactivity (González *et al*., 2016).

### Moss photosynthetic capacity

This study shows that *Sphagnum* mosses can maintain photosynthetic activity under waterlogged conditions, but their performance varies markedly among species and is strongly influenced by the pH of the surrounding solution. The pronounced interspecific and pH-dependent differences observed among the five species examined (*S. auriculatum*, *S. palustre*, *S. balticum*, *S. girgensohnii*, and *S. strictum*), are in line with previous findings that ecological specialization of *Sphagnum* spp. is tightly linked to the environmental pH (Harley *et al*., 1989; Haraguchi, 1996; Van Gaalen *et al*., 2007). These differences had impact on CO_2_ assimilation, light-use efficiency, photoprotective capacity and respiration, and hence presumably on the adaptability of these species to variable environmental conditions. For example, *S. strictum* displayed the highest A_max_ at pH 3, indicating a potentially high physiological adaptation to strong acidic conditions. In contrast, *S. auriculatum* performed best at more alkaline conditions, while *S. palustre* maintained stable *Aₘₐₓ* values across the entire pH range. *Sphagnum* mosses are generally known to grow better in wet, acidic conditions, and be negatively impacted by alkaline waters (Koks *et al*., 2019). However, *S. auriculatum* is reported also as moderately acidophilic (Hugonnot & Chavoutier, 2024), while *S. palustre* is a species occurring in regions with HCO_3_^−^ rich surface water, cultivated for paludiculture (Koks *et al*., 2025), suggesting its potential ecological advantages as a pH-generalist in heterogeneous peatland environments. *Sphagnum girgensohnii* exhibited maximum photosynthetic performance at pH 5. This pattern contrasts with reports by Haraguchi *et al*. (2003) who found a photosynthetic maximum for *S. girgensohnii* around pH 7.2, although the authors noted that its variation across pHs was minimal. In contrast, *S. balticum* exhibited consistently low assimilation rates from acidic to neutral pH levels, indicating limited competitive ability and sensitivity to chemical fluctuations, compared to the other species.

Light-response curves for the five investigated *Sphagnum* species showed that the saturation of photosynthesis occurred between 100 and 300 μmol m⁻² s⁻¹ PAR, consistent with typical adaptation to shade/low-light conditions of *Sphagnum* mosses (Harley *et al*., 1989; Davey & Rothery, 1997; Van Gaalen *et al*., 2007; Bonnet *et al*., 2010). This adaptation varies among species and pHs. For example, Maseyk *et al*. (1999) reported a light saturation in *S. cristatum* and *S. australe* from New Zealand ranging from 111 to 266 μmol m⁻² s⁻¹, whereas *S. angustifolium* from Alaska exhibited saturation between 250 and 500 μmol m⁻² s⁻¹, with an LCP around 37 μmol m⁻² s⁻¹ (Harley *et al*., 1989). In our study, *S. girgensohnii* showed the highest photosynthesis performance and the lowest LCP (34 μmol m⁻² s⁻¹) at pH 5, reflecting efficient carbon assimilation in low-light conditions of its natural habitat (Laine *et al*., 2018). Conversely, *S. balticum* exhibited much higher LCP values, especially at pH 5 and 7 (∼124 μmol m⁻² s⁻¹), suggesting a stronger dependence on high irradiance to achieve positive carbon balance, in agreement with the low-light limitations reported for acid-sensitive *Sphagnum* species (Van Gaalen *et al*., 2007).

The non-photochemical quenching (NPQ) is a photoprotective mechanism of plant photosystems that dissipates excess light energy as heat, generally increasing with light intensity (Van Gaalen *et al*., 2007; Lu *et al*., 2022;). The NPQ patterns exhibited by the five studied *Sphagnum* spp. were not always consistent with CO_2_ assimilation trends. For example, *S. strictum*, despite its A_max_ peak at pH 3, exhibited maximum NPQ values at pH 7 and 9, suggesting enhanced photoprotective energy dissipation under suboptimal chemical conditions to mitigate photooxidative stress (Van Gaalen *et al*., 2007). In contrast, *S. auriculatum* and *S. girgensohnii* showed gradual NPQ responses across pH levels, indicating efficient energy regulation, while low variation in NPQ for *S. palustre* despite pH treatments, as well as in CO_2_ assimilation, corroborated its status as stress-tolerant and physiologically versatile species (Keightley *et al*., 2024).

The photochemical efficiency of PSII, expressed through dark-adapted (F_v_/F_m_) and light-adapted (F_v_’/F_m_’) fluorescence parameters, further reflected species-specific responses to pH and light conditions. The F_v_’/F_m_’ declined with increasing PAR in all species with a steeper decrease within the first 200 μmol m⁻² s⁻¹, consistent with the typical downregulation of PSII under excess light, as expected for shade-adapted mosses (Van Gaalen *et al*., 2007; Sepp & Ilomets, 2008). Differences in dark-acclimated F_v_/F_m_ among species were minor, apart from *S. auriculatum*, suggesting limited pH effects on PSII photochemistry in darkness. Notably, *S. strictum* maintained relatively low F_v_/F_m_ values, along with high NPQ at neutral pH, indicative of sustained engagement of photoprotective pathways and potential chronic photoinhibition. Conversely, *S. palustre* and *S. girgensohnii* maintained higher F_v_/F_m_ under alkaline conditions, reflecting efficient PSII repair and recovery. The combination of moderate NPQ and high F_v_/F_m_ values indicates efficient balance between photon use and energy dissipation, which allow continuous carbon assimilation under variable environmental stress (Korrensalo *et al*., 2017).

The quantum efficiency of PSII (*Φ*) was highest in *S. strictum* across all pHs, despite reduced PSII efficiency, suggesting a trade-off between photochemical performance and protective energy dissipation. *Sphagnum girgensohnii* showed lower *Φ* coupled with decreased A_max_ at pH 3 and 7, while other species showed relatively invariant average *Φ* values across pH treatments, highlighting divergent energy management strategies (Van Gaalen *et al*., 2007). Dark respiration rates (R_d_), indicating the rate at which the mosses release CO_2_ in the absence of light (negative values), varied significantly, with *S. strictum* and *S. balticum* exhibiting higher R_d_, implying elevated metabolic costs or stress-related carbon loss that limits net assimilation (Robroek *et al*., 2009). The combination of low A_max_ and high R_d_ in *S. balticum*, except in alkaline solution, highlights its poor photosynthetic competitiveness observed under most pH conditions.

Overall, our findings suggest that among the 5 *Sphagnum* species, *S. palustre*, beyond functioning as efficient ion adsorbent, also emerged as the most stress-tolerant and generalist species, maintaining stable CO_2_ assimilation, efficient PSII function, and limited photoprotective upregulation across all tested pHs. In contrast, *S. auriculatum*, *S. girgenshonii*, and S. *strictum* performed well only under narrow pH ranges, while *S. balticum* showed poor competitiveness across most treatments.

### Moss chemical reactivity and photosynthetic capacity

In *Sphagnum* spp., net photosynthesis increases with rising tissue water content until an optimum is reached, beyond which rates decline (Jauhiainen & Silvola, 1999). Under submerged or water-saturated conditions, photosynthesis is often limited by slow CO_2_ diffusion in water (Murray *et al*., 1989; Silvola, 1990; Li *et al*., 1992; Williams & Flanagan, 1996; Van Gaalen *et al*., 2007) compared with air (Massman, 1998). CO_2_ uptake therefore depends on dissolved inorganic carbon availability. In water, CO_2(g)_ dissolves forming H_2_CO_3(aq)_, which dissociates into H⁺_(aq)_ and HCO_3_⁻_(aq)_. pH shifts these equilibria, with higher pH driving carbon toward bicarbonate and carbonate, thereby reducing free CO_2(g)_ (*e.g.,* Allen & Spence, 1981; Raven *et al*., 1998).

Species with low pH_pzc_ values, such as S. *angustifolium* (4.4), *S. medium*, and S. *teres* (4.6) remain negatively charged in a broader pH range, enhancing proton exchange and acidification of the boundary water layer, thereby shifting local equilibria toward CO_2(g)_ potentially favour photosynthesis. Conversely, species with high pH_pzc_ like *S*. *strictum* (8.2), *S. subnitens* (6.9) and *S*. *girgensohnii* (6.8) could be less efficient in acidifying their surroundings under comparable environmental conditions, potentially leading to lower passive CO_2_ uptake in neutral-to-alkaline waters. Among the five species analysed for photosynthesis, *S. palustre* and *S. balticum* showed the lowest pH_pzc_, and hence a potentially highest acidification capacity and photosynthetic rates across a broader pH range in natural environments. This effect should be most pronounced at pH values above pH_pzc_, *i.e.,* at pH values > 6.3, 5.3, 6.8, 4.9 and 8.2 respectively for *S. auriculatum*, *S. balticum*, *S. girgensohnii*, *S. palustre*, and *S. strictum*. Except for *S. palustre*, which maintained relatively stable maximum photosynthetic rates across all tested pH levels, indicating a broad tolerance, *S. auriculatum* and *S. balticum* maximum CO_2_ assimilation occurred at pH 9, well above their pH_pzc_, whereas photosynthetic rates at pH 3–7 remained comparatively low. By contrast, *S. girgensohnii* exhibited its highest assimilation at pH 5, below its pH_pzc_, and strongly reduced rates at both more acidic and more alkaline conditions. *Sphagnum strictum* reached its highest assimilation at pH 3, far below its pH_pzc_, and showed only moderate to low rates at higher pH values. These findings suggest that pH_pzc_ alone is not a reliable predictor of *Sphagnum* photosynthetic capacity.

Overall, the chemical reactivity of *Sphagnum* surfaces can predict photosynthetic capacity to a certain extent, when considering specific functional groups and pH. As supported by correlations (Supporting Information Table **S2**) between binding-site abundance and light-response parameters (A_max_, R_d_, LCP and *Φ*), functional groups providing high proton adsorption and negative surface charge more likely regulate micro-scale pH and CO_2_ speciation in the water film surrounding *Sphagnum* surfaces. Carboxyls acidify the immediate boundary layer between moss surfaces and the water film, driving the conversion of bicarbonate to free CO_2_ at the cell surface. This localized acidification, particularly within hyalocysts and around photosynthetically active tissues, can enhance CO_2_ solubility and promote its passive diffusion, ultimately shifting the chemical equilibrium toward greater CO_2_ uptake (Glime, 2020). While carboxyls mainly act as proton donors, phosphodiesters help stabilize negative surface charge and bind cations (Ca²⁺, Mg²⁺, H⁺), thereby retaining water and maintaining the proton-enriched zones generated by carboxyls (Clymo, 1964; Vijayaraghavan & Yun, 2008). Such microzones arise from stabilized protonated states and shifts in interfacial *pK_a_* (Ottosson *et al*., 2011; Gerland *et al*., 2020), influencing proton-coupled processes at plant surfaces. Phosphoryl groups can also bind and release protons, regulating micro-scale pH (Baker *et al*., 2009). As polyprotic acids with higher negative charge density than carboxyls (Franz, 2001), they may more effectively enhance CO_2_ assimilation under bicarbonate-rich conditions. This is consistent with stronger correlations between A_max_ and phosphoryls amount compared to carboxyls (Supporting Information Table **S2**). Phosphoryls also correlate strongly with CO_2_ fluxes from dark respiration, suggesting that their proton- and cation-binding properties help retain CO_2_ released by respiration in the boundary layer, increasing its potential for reassimilation. In contrast, amine groups correlate weakly with *Aₘₐₓ* at neutral–alkaline pH (Supporting Information Table **S2**). Near neutral pH, amines occur mainly as –NH_3_^+^ and act as basic sites that can alkalinize the microenvironment, shifting CO_2_ toward bicarbonate and carbonate, and thus reducing free CO_2_ availability (Jiang *et al*., 2004; Hamdy *et al*., 2021).

The LCP reflects the irradiance at which photosynthesis balances respiration. Surfaces that promote CO_2_ enrichment (via carboxyls, phosphoryls, phosphodiesters) should enable positive net photosynthesis at lower light intensities, thus lowering the LCP. Overall, functional group composition favours higher photosynthetic efficiency under neutral to alkaline pH conditions, with phosphoryls acting as the strongest promoters across all conditions.

Carboxyls show weak correlations with A_max_ but consistent links to LCP at low pH (Supporting Information Table **S2**), suggesting that although they increase CO_2_ availability at high pH, excessive proton release may increase stress under acidic conditions. Amine-rich surfaces show a positive correlation with R_d_ and LCP (Supporting Information Table **S2**), indicating reduced productivity under shaded or low-light conditions. Amine-rich surfaces correlate positively with R_d_ and LCP, indicating lower productivity at low irradiance; their negative influence becomes clearest at pH ∼9, where they increase LCP. Quantum yield (*Φ*), reflecting CO_2_ fixation per absorbed photon, should depend strongly on CO_2_ availability at low light. Although the alkalinizing effect of amines would predict a negative impact on *Φ*, correlations instead show that amines are the most consistent positive contributors across nearly all pH levels (*r* = 0.74–0.94). This paradox suggests that, despite reducing free CO_2_, amines may enhance photon-use efficiency through microenvironmental charge stabilization or support of bicarbonate-use pathways.

It is important to note that acid–base titrations reflect only passive biosurface properties, as samples were devitalized prior to analysis, whereas photosynthesis measurements involve living tissues, and thus integrate both passive and active mechanisms. Despite methodological differences across studies, assimilation rates observed here align with published values (Haraguchi *et al*., 2003; Bengtsson *et al*., 2016; Jassey & Signarbieux, 2019), indicating that waterlogging did not impose significant limitations.

Beyond the buffering capacity and chemical reactivity of moss surfaces, additional mechanisms likely contribute to sustaining photosynthesis under aquatic conditions.

The effects of water content on photosynthesis rate can greatly depend on the moss structure (Silvola, 1991). Rice & Schuepp (1995) demonstrated that *Sphagnum* traits, such as leaf size and the degree of chlorocystes exposure at leaf surface, can suppress the CO_2_ diffusion in surrounding water layers. Conversely, aquatic taxa with thinner branch leaves, loosely arranged foliage, and highly exposed photosynthetic cells lower overall resistance to CO_2_ uptake. In our study, the aquatic/hollow species *S. cuspidatum*, *S. balticum*, and *S. tenellum*, despite a surface chemistry allowing acidification of boundary water layer, generally perform less efficiently than non-aquatic species under neutral to alkaline conditions. Under acidic conditions, however, their performance equals or exceeds that of non-aquatic species. *Sphagnum aongstroemii*, typically from wet but not strictly aquatic hollows (Supporting Information Table **S1**), appears more physiologically versatile than *S. cuspidatum*, *S. balticum*, and *S. tenellum*. The occurrence of bryophytes in aquatic habitats with pH >7 indicates an ability to acquire CO_2_ even under alkaline conditions (Glime & Vitt, 1984). Thin wax layers may protect leaves from waterlogging and facilitate CO_2_ diffusion in saturated environments (Proctor, 1984), while the empty hyalocysts and leaf–stem architecture may trap air bubbles, providing additional CO_2_ sources. Other mechanisms observed in aquatic mosses, such as CO_2_-concentrating strategies in *Fontinalis antipyretica* and *Fissidens* spp., or the external use of carbonic anhydrase, may also apply to *Sphagnum* in facilitating CO_2_ availability, even though there is still no evidence of direct use of HCO_3_⁻ as carbon source (Peñuelas, 1985; Raven *et al*., 1998).

Cultivation of *S. centrale*, *S. cuspidatum*, *S. papillosum*, and *S. squarrosum* in bioreactors in liquid medium, although the sterile and uniform environment may limit plasticity in response to stressors (*e.g.,* fluctuating light, pH, and nutrient availability, microbial interactions), preserved the core functional groups, the ion exchange properties and pH regulation ability of field-grown conspecific mosses, as previously demonstrated (González *et al*., 2016; Di Palma *et al*., 2019). This retained biosurface chemistry, as well as the morphological features (Beike *et al*., 2015) suggests that the photosynthetic performance of bioreactor-grown clones is largely maintained. This interpretation is supported by recent work showing that micropropagated clones exhibited efficient CO_2_ assimilation and photochemical efficiency, despite the absence of natural environmental variability (Keightley *et al*., 2024). Furthermore, the optimal nutrient availability and reduced stress in bioreactors could minimize photoinhibition and metabolic disruption, allowing the full expression of photosynthetic capacity without the constraints typically imposed in field settings (Schipperges & Rydin, 1998; Keightley *et al*., 2024). Overall, the homogeneity among clones in terms of surface chemistry and growth conditions highlights the potential of controlled cultivation to reproduce key physicochemical properties of natural populations, providing predictable and reproducible physiological performance. This supports the use of *Sphagnum* clones as model organisms for experimental research and for large-scale applications, including wastewater treatment, biomonitoring, and peatland restoration.

## Acknowledgments

This study was funded by the National Recovery and Resilience Plan (NRRP), Mission 4 Component 2 Investment 1.4 - Call for tender No. 3138 of 16 December 2021, rectified by Decree n.3175 of 18 December 2021 of Italian Ministry of University and Research funded by the European Union – NextGenerationEU, Award Number: Project code CN_00000033, Concession Decree No. 1034 of 17 June 2022 adopted by the Italian Ministry of University and Research, CUP B83C22002930006, Project title “National Biodiversity Future Center - NBFC”. RR acknowledges financial support by the German Federal Ministry of Food and Agriculture BMEL (MOOSzucht, No. 22007216, MOOSstart, No. 2221MT020B), the Lower Saxony Ministry of Food, Agriculture and Consumer Protection (PALUDIFarming, No. 105-29213-2940/2022), and by Deutsche Forschungsgemeinschaft (DFG, German Research Foundation) under Germany’s Excellence Strategy (*liv*MatS - EXC-2193/1 – Project ID 390951807 and CIBSS – EXC-2189 – Project ID 390939984).

We thank Jukka Laine, Prof. Emeritus at the University of Helsinki, for moss field collection and photography, Dr. Pierluigi Cox for greenhouse cultivation of mosses after all analytical determinations, and Britta Rothgänger for the images of the *Sphagnum* clones.

## Author Contributions

ADP: Leading conceptualization and design of the work, experimental work, formal analysis, data curation, writing, review and editing the original draft; EP: support in photosynthesis experiment implementation, and gas exchange data curation and discussion; AGG and OSP: support in chemical data analysis and interpretation; RR: cultivation of mosses in photobioreactors and review of original draft; JG: review of original draft and language editing; NK: support in chemicaldata interpretation; CC: supervision and funding acquisition. All authors contributed to review the original draft and have approved the final manuscript.

## Supporting Information

**Fig. S1.**
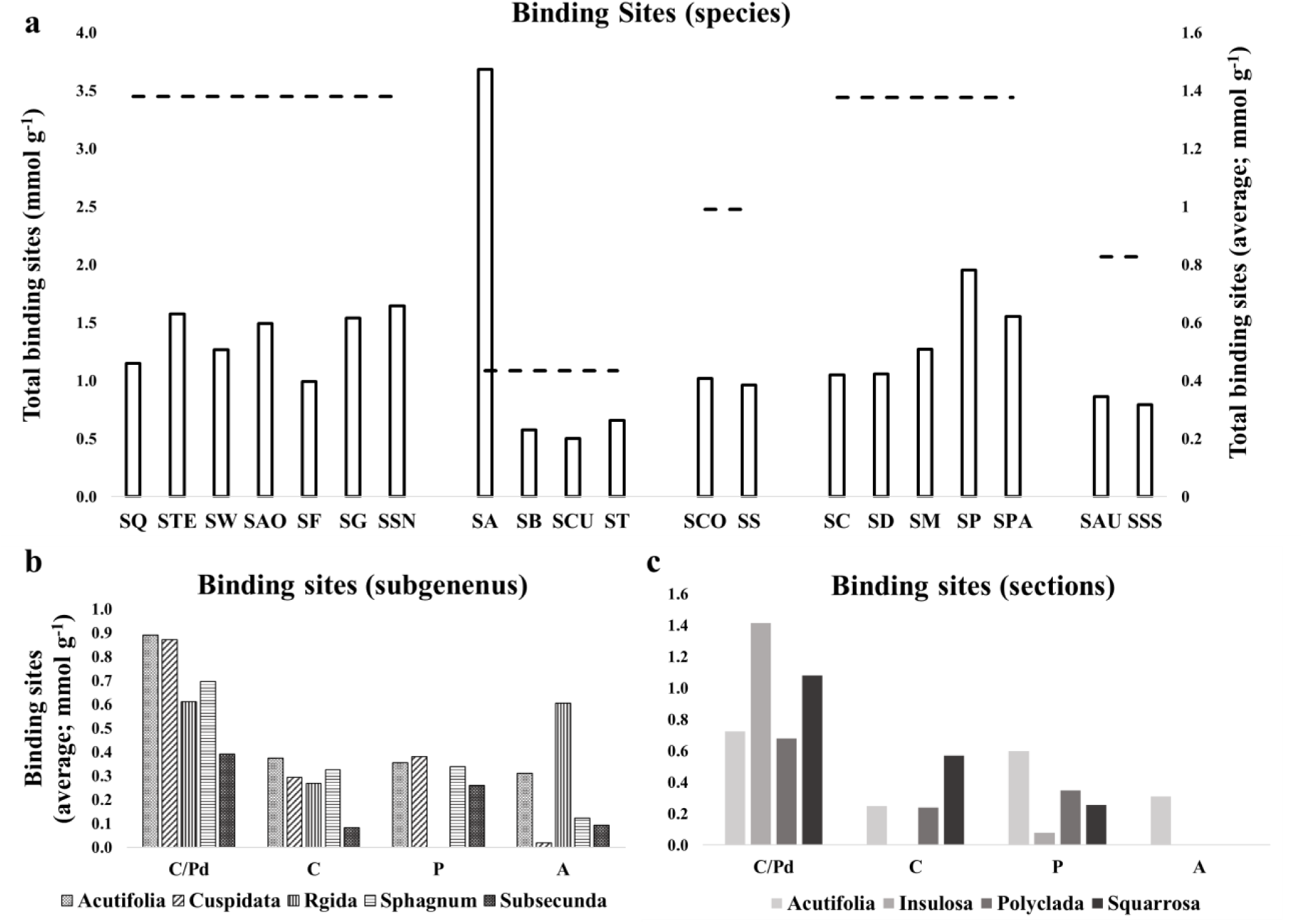
Amount of binding sites (mmol g^-1^; *n* = 3) of the 20 *Sphagnum* species collected from nature, computed from acid-base titration curves by applying LPM model. Data are shown as total values and for each type of binding site (C = carboxyl; Pd =phosphodiester; P = phosphoryl; A = amine), and distinguished for moss species (a), subgenus (b) and section (c). SA = *S. angustifolium*; SAO = *S. aongstroemii*; SAU = *S. auriculatum*; SB = *S. balticum*; SC = *S. centrale*; SCO = *S. compactum*; SD = *S. divinum*; SF = *S. fimbriatum*; SG = *S. girgensohnii*; SM = *S. medium*; SP = *S. palustre*; SPA = *S. papillosum*; SQ = *S. squarrosum*; SS = *S. strictum*; SSN = *S. subnitens*; SSS = *S. subsecundum*; SCU = *S. cuspidatum*; ST = *S. tenellum*; STE = *S. teres*; SW = *S. wulfianum*.

**Fig. S2.**
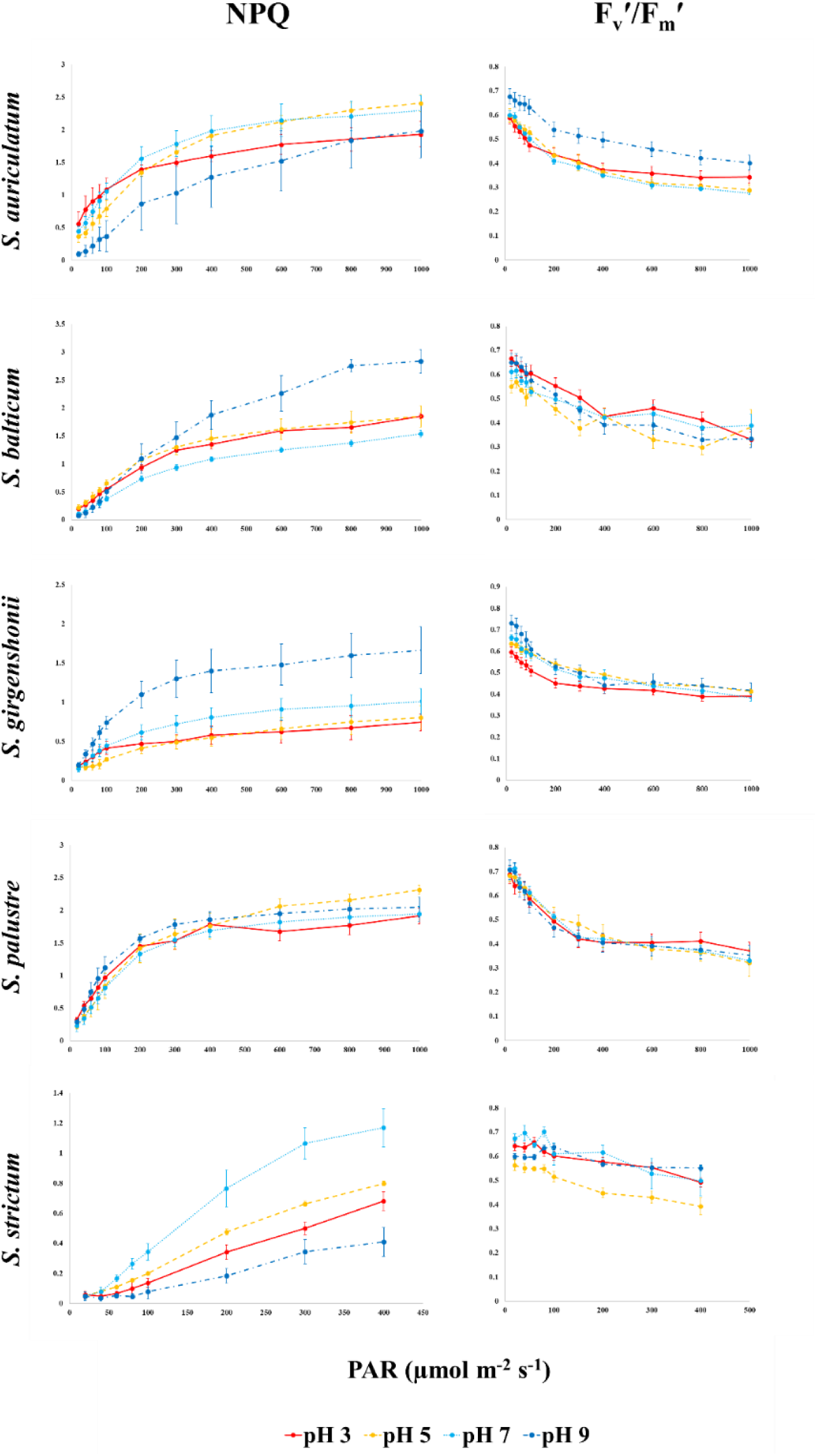
Curves of NPQ (Non Photochemical Quenching) and F_v_’/F_m_’ (maximum efficiency of PSII photochemistry in the light-adapted state) as function of PAR (Photosynthetically Active Radiation; µmol m^-2^ s^-1^), for 5 *Sphagnum* species collected in the field and four pH values (*n* = 5).

**Table S1.**
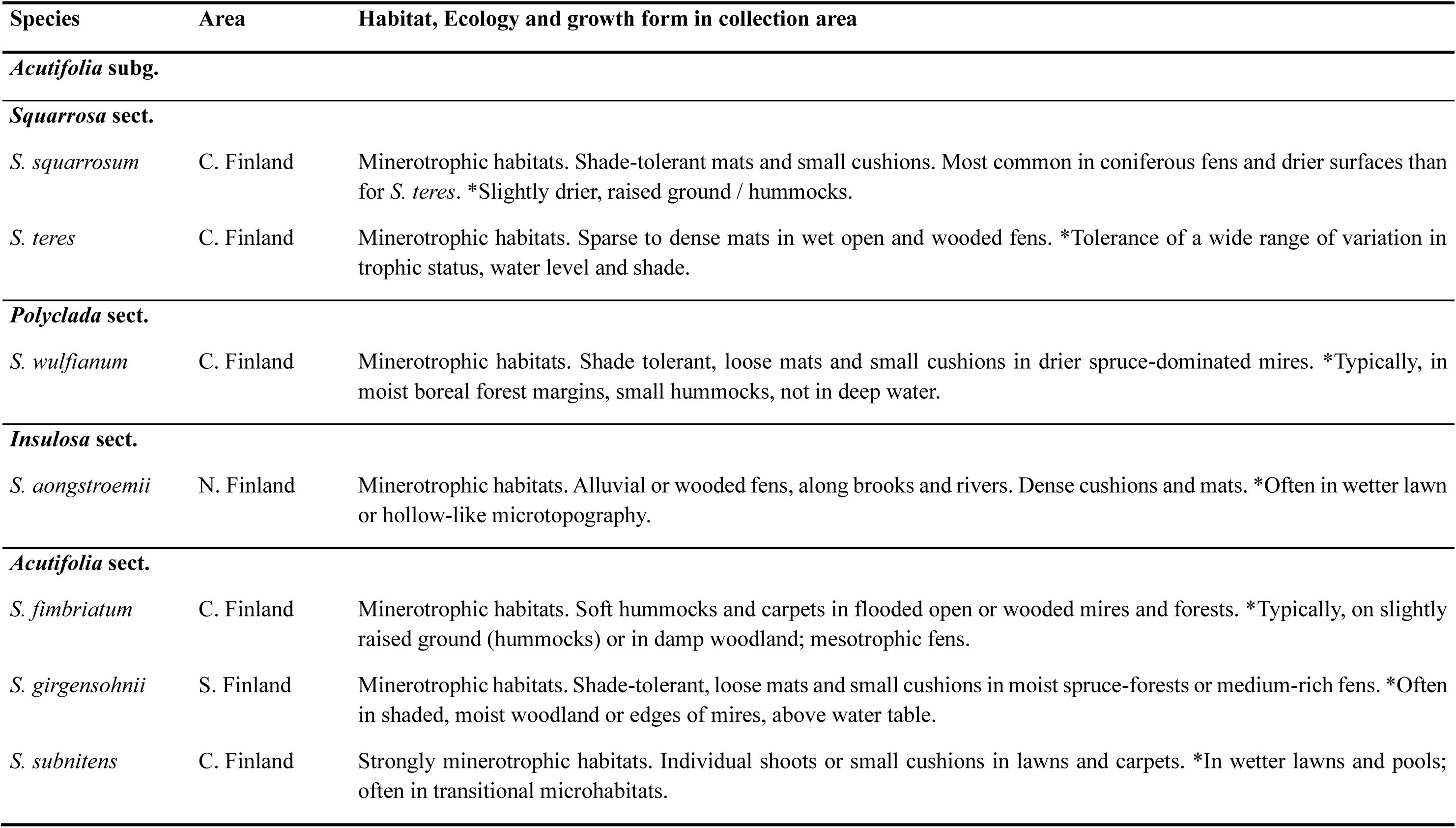
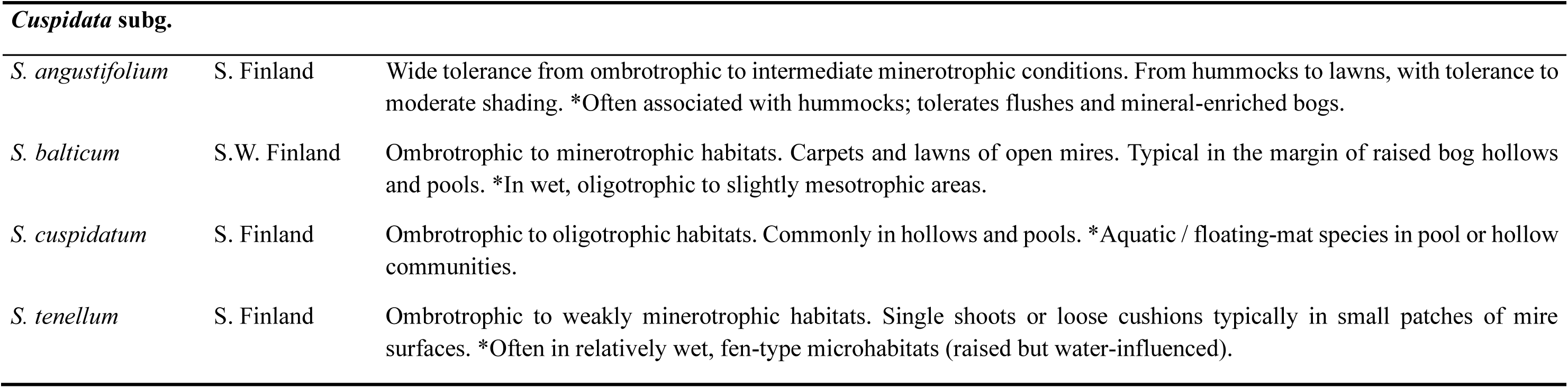
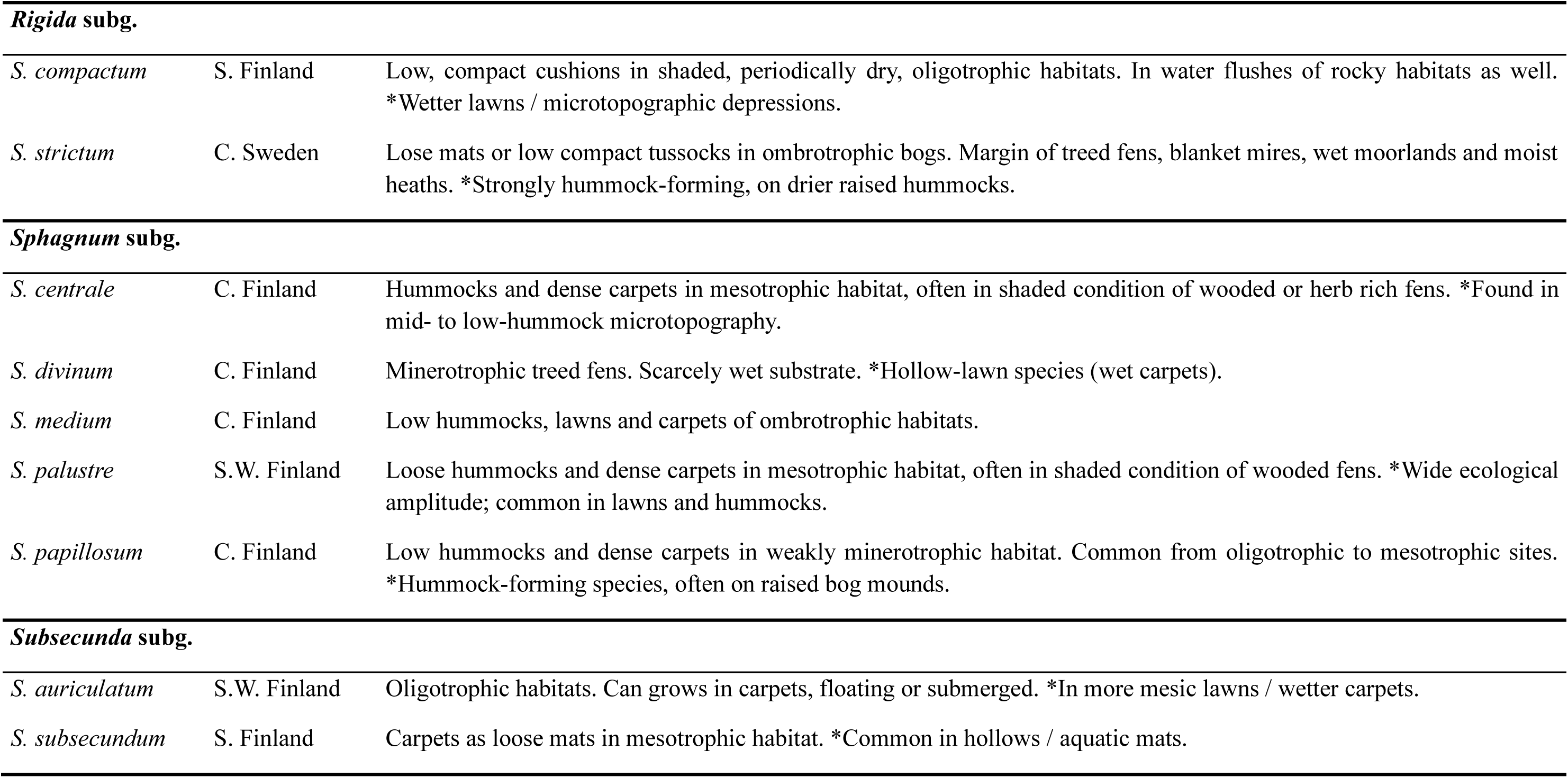
*Sphagnum* species used for the experiments, along with the collection areas, habitats, and growth form (Laine *et al*., 2018). *Other descriptions from Daniels & Eddy (1985). C. = central; S. = southern; N. = northern; W. =western.

| Species | Area | Habitat, Ecology and growth form in collection area |
| --- | --- | --- |
| *Acutifolia* subg. |  |  |
| *Squarrosa* sect. |  |  |
| *S. squarrosum* | C. Finland | Minerotrophic habitats. Shade-tolerant mats and small cushions. Most common in coniferous fens and drier surfaces than for *S. teres*. *Slightly drier, raised ground / hummocks. |
| *S. teres* | C. Finland | Minerotrophic habitats. Sparse to dense mats in wet open and wooded fens. *Tolerance of a wide range of variation in trophic status, water level and shade. |
| *Polyclada* sect. |  |  |
| *S. wulfianum* | C. Finland | Minerotrophic habitats. Shade tolerant, loose mats and small cushions in drier spruce-dominated mires. *Typically, in moist boreal forest margins, small hummocks, not in deep water. |
| *Insulosa* sect. |  |  |
| *S. aongstroemii* | N. Finland | Minerotrophic habitats. Alluvial or wooded fens, along brooks and rivers. Dense cushions and mats. *Often in wetter lawn or hollow-like microtopography. |
| *Acutifolia* sect. |  |  |
| *S. fimbriatum* | C. Finland | Minerotrophic habitats. Soft hummocks and carpets in flooded open or wooded mires and forests. *Typically, on slightly raised ground (hummocks) or in damp woodland; mesotrophic fens. |
| *S. girgensohnii* | S. Finland | Minerotrophic habitats. Shade-tolerant, loose mats and small cushions in moist spruce-forests or medium-rich fens. *Often in shaded, moist woodland or edges of mires, above water table. |
| *S. subnitens* | C. Finland | Strongly minerotrophic habitats. Individual shoots or small cushions in lawns and carpets. *In wetter lawns and pools; often in transitional microhabitats. |
| ***Cuspidata* subg.** |  |  |
| *S. angustifolium* | S. Finland | Wide tolerance from ombrotrophic to intermediate minerotrophic conditions. From hummocks to lawns, with tolerance to moderate shading. *Often associated with hummocks; tolerates flushes and mineral-enriched bogs. |
| *S. balticum* | S.W. Finland | Ombrotrophic to minerotrophic habitats. Carpets and lawns of open mires. Typical in the margin of raised bog hollows and pools. *In wet, oligotrophic to slightly mesotrophic areas. |
| *S. cuspidatum* | S. Finland | Ombrotrophic to oligotrophic habitats. Commonly in hollows and pools. *Aquatic / floating-mat species in pool or hollow communities. |
| *S. tenellum* | S. Finland | Ombrotrophic to weakly minerotrophic habitats. Single shoots or loose cushions typically in small patches of mire surfaces. *Often in relatively wet, fen-type microhabitats (raised but water-influenced). |

**Table S1.** Continued.
| Species | Area | Habitat, Ecology and growth form in collection area |
| --- | --- | --- |
| ***Rigida* subg.** |  |  |
| *S. compactum* | S. Finland | Low, compact cushions in shaded, periodically dry, oligotrophic habitats. In water flushes of rocky habitats as well. *Wetter lawns / microtopographic depressions. |
| *S. strictum* | C. Sweden | Lose mats or low compact tussocks in ombrotrophic bogs. Margin of treed fens, blanket mires, wet moorlands and moist heaths. *Strongly hummock-forming, on drier raised hummocks. |
| ***Sphagnum* subg.** |  |  |
| *S. centrale* | C. Finland | Hummocks and dense carpets in mesotrophic habitat, often in shaded condition of wooded or herb rich fens. *Found in mid- to low-hummock microtopography. |
| *S. divinum* | C. Finland | Minerotrophic treed fens. Scarcely wet substrate. *Hollow-lawn species (wet carpets). |
| *S. medium* | C. Finland | Low hummocks, lawns and carpets of ombrotrophic habitats. |
| *S. palustre* | S.W. Finland | Loose hummocks and dense carpets in mesotrophic habitat, often in shaded condition of wooded fens. *Wide ecological amplitude; common in lawns and hummocks. |
| *S. papillosum* | C. Finland | Low hummocks and dense carpets in weakly minerotrophic habitat. Common from oligotrophic to mesotrophic sites. *Hummock-forming species, often on raised bog mounds. |
| ***Subsecunda* subg.** |  |  |
| *S. auriculatum* | S.W. Finland | Oligotrophic habitats. Can grows in carpets, floating or submerged. *In more mesic lawns / wetter carpets. |
| *S. subsecundum* | S. Finland | Carpets as loose mats in mesotrophic habitat. *Common in hollows / aquatic mats. |

**Table S2.** Pearson correlations between amounts of binding sites (total and by type) and photosynthesis parameters. BS = binding sites; C-Pd = carboxyls-phosphodiesters; C= carboxyls; P = phosphoryls; A = amines; *r* = correlation coefficient; *r^2^*= coefficient of determination; A_max_ = maximum photosynthetic rate; *Φ* 4 = Apparent quantum yield; LCP = Light compensation point; R_d_ = dark respiration.

|  | pH | Total BS |  | C-Pd |  | C |  | P |  | A |  |
| --- | --- | --- | --- | --- | --- | --- | --- | --- | --- | --- | --- |
| | | $r$ | $r^2$ | $r$ | $r^2$ | $r$ | $r^2$ | $r$ | $r^2$ | $r$ | $r^2$ |
| $A_{\max}$ | 3 | 0.14 | 0.02 | 0.74 | 0.54 | 0.01 | 0.00 | -0.21 | 0.04 | 0.76 | 0.57 |
|  | 5 | 0.69 | 0.47 | 0.78 | 0.61 | -0.02 | 0.00 | 0.57 | 0.33 | 0.46 | 0.21 |
|  | 7 | 0.72 | 0.52 | 0.78 | 0.60 | -0.17 | 0.03 | 0.67 | 0.44 | 0.11 | 0.01 |
|  | 9 | -0.49 | 0.24 | -0.42 | 0.18 | -0.73 | 0.53 | -0.16 | 0.02 | 0.31 | 0.10 |
| $\Phi$ | 3 | -0.13 | 0.02 | 0.14 | 0.02 | 0.13 | 0.02 | -0.55 | 0.30 | 0.74 | 0.54 |
|  | 5 | -0.02 | 0.00 | 0.73 | 0.53 | 0.04 | 0.00 | -0.37 | 0.14 | 0.94 | 0.89 |
|  | 7 | -0.07 | 0.01 | 0.07 | 0.00 | 0.13 | 0.02 | -0.45 | 0.21 | 0.41 | 0.17 |
|  | 9 | -0.19 | 0.04 | 0.07 | 0.00 | 0.04 | 0.00 | -0.50 | 0.25 | 0.94 | 0.89 |
| LCP | 3 | -0.01 | 0.00 | -0.06 | 0.00 | 0.83 | 0.68 | -0.78 | 0.61 | 0.15 | 0.02 |
|  | 5 | -0.46 | 0.21 | -0.48 | 0.23 | 0.46 | 0.22 | -0.73 | 0.53 | -0.01 | 0.00 |
|  | 7 | -0.65 | 0.42 | -0.66 | 0.44 | 0.14 | 0.02 | -0.88 | 0.78 | 0.05 | 0.00 |
|  | 9 | -0.42 | 0.17 | -0.62 | 0.39 | -0.10 | 0.01 | -0.71 | 0.50 | 0.66 | 0.44 |
| $R_d$ | 3 | 0.42 | 0.18 | 0.52 | 0.27 | -0.31 | 0.10 | 0.85 | 0.72 | -0.71 | 0.50 |
|  | 5 | 0.55 | 0.31 | 0.77 | 0.60 | -0.11 | 0.01 | 0.89 | 0.80 | -0.63 | 0.39 |
|  | 7 | 0.49 | 0.24 | 0.52 | 0.27 | -0.15 | 0.02 | 0.87 | 0.76 | -0.36 | 0.13 |
|  | 9 | 0.29 | 0.09 | 0.60 | 0.36 | 0.07 | 0.00 | 0.57 | 0.33 | -0.83 | 0.69 |

## Methods S1

LPM uses a grid of fixed p*K* values and optimizes the binding site concentrations. In each case, zero is considered as a possible solution then, a p*K* spectrum is produced for the discrete binding sites computed by the number of p*K* values. The acid-base titration experiments can be modeled according with the proton dissociation reaction for a single protonated site:

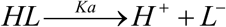

where *HL* are the protonated binding sites on the moss surface and *H^+^* is the hydrogen concentration measured directly with a pH electrode. *L^-^* is the deprotonated surface reactive site with a net negative charge. Accordingly, the apparent proton dissociation constant is:

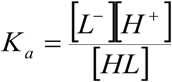

where p*K_a_*=-log_10_*K_a_*. Therefore, the experimental net surface charge excess for the *i*th addition of titrant (*b_meas,i_*) is:

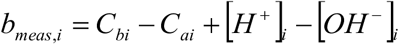

where *C_ai_* and *C_bi_* are the acid base concentrations at the *i*th addition of titrant. The calculated net surface charge excess can be written as:

[umath4

Here, *m* is the number of binding sites and [*L_T_*] is the site density. *K_a,j_* is the apparent acidity constant for the *j*th site, and *S_0_* is a constant which is necessary in order to account for positive changes on the surface. In the case of pH-edge adsorption, within the formalism of *K_s,j_*, the reaction between a metal in solution and available binding sites on the moss cell surface is defined as:

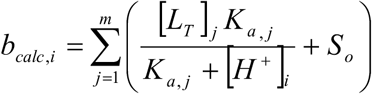

where *Me^2+^* represents a generic metal divalent cation, *B_j_* is a surface reactive site and *K_s,j_* is the apparent concentration equilibrium constant conditional on ionic strength. For a *j-*th deprotonated binding sites at the *i-*th pH values, *K_s,j_* is:

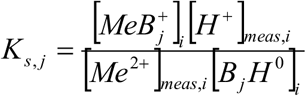

where *K_s,j_* is a function of experimentally measured proton and metal concentrations ([H^+^]*_meas,i_* and [*Me^2^*^+^]*_meas,i_*) and the amount of Me^2+^ bound to the *j-*th site at the *i-*th pH value ([*MeB* ^+^] ). In the case of Langmuirian isotherm adsorption (fixed pH), the LPM considers the reaction between metal and binding sites expressed as:

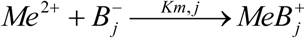

where *B_j_* is the specific surface functional group and *K_m,j_* is the apparent metal-ligand binding constant conditional on ionic strength, for a *j-*th deprotonated functional group at fixed pH value is written as:

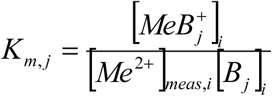

*K_m,j_* is a function of the experimental metal concentration ([*Me^2+^*]*_meas,i_*) and of the amount of Me^2+^ bound to the *j-*th site as a function of increasing biomass and at fixed pH value ([*MeB ^+^*] ). As described for pH-edge data, the available binding sites on moss surfaces are computed and assigned to a fixed p*K_m,j_* grid. In addition, the values of *K_s,j_* and *K_m,j_* computed cannot be directly comparable because K_s_ is a function of *K_m_* (*K_s_*= *K_m_*·*K_a_*), where *K_a_* is the acidity constant for a specific functional group on the moss surface.

